# Population genomics reveals demographic history and climate adaptation in Japanese *Arabidopsis halleri*

**DOI:** 10.1101/2024.04.08.588504

**Authors:** Ryo A Suda, Shosei Kubota, Vinod Kumar, Vincent Castric, Ute Krämer, Shin-Ichi Morinaga, Takashi Tsuchimatsu

## Abstract

Climate oscillations in the Quaternary forced species to major latitudinal or altitudinal range shifts. It has been suggested that adaptation concomitant with range shifts plays key roles in species responses during climate oscillations, but the role of selection for local adaptation to climatic changes remains largely unexplored. Here, we investigated population structure, demographic history, and signatures of climate-driven selection based on genome-wide polymorphism data of 141 Japanese *Arabidopsis halleri* individuals, with European ones as outgroups. Coalescent-based analyses suggested a genetic differentiation between Japanese subpopulations since the Last Glacial Period (LGP), which would have contributed to shaping the current pattern of population structure. Population demographic analysis revealed the population size fluctuations in the LGP, which were particularly prominent since the subpopulations started to diverge (∼50 kya). The ecological niche modeling predicted the range shifts from southern coastal regions to northern coastal and mountainous areas, possibly in association with the population size fluctuations. Through genome-wide association analyses of bioclimatic variables and selection scans, we investigated whether climate-associated loci are enriched in the extreme tails of selection scans, and demonstrated the prevailing signatures of selection, particularly toward a warmer climate in southern subpopulations and a drier environment in northern subpopulations, which may have taken place during or after the LGP. Our study highlights the importance of integrating climate associations, selection scans, and population demographic analyses for identifying genomic signatures of population-specific adaptation, which would also help us predict the evolutionary responses to future climate changes.

## Introduction

Climate oscillations in the Quaternary (2.6 Mya to present) forced species to major latitudinal or altitudinal range shifts (Comes and Kadereit, 1998; Hewitt, 2004; Qiu et al., 2011; Schmitt, 2007). In glacial periods, some locations served as climate refugia, where species were allowed to persist in local habitats, and they expanded from there when climate conditions improved (Gavin et al., 2014; Provan and Bennett, 2008). The repeated environmental changes would lead to range shifts, contractions, expansions, and subdivisions of populations that profoundly shaped the present-day distributions and genetic structures of many plant and animal species (Hewitt, 2000, 2004).

It has been suggested that adaptation concomitant with range shifts plays key roles in species responses during climate oscillations in the Quaternary (Davis and Shaw, 2001; De Lafontaine et al., 2018; Excoffier et al., 2009; Keller et al., 2011; Luqman et al., 2023). Understanding the mechanisms of local adaptation to past climate changes is also crucial for predicting the evolutionary responses to future environmental changes (Waldvogel et al., 2020). However, the influence of past climate oscillations has mainly been studied through the lens of neutral genetic variation, and knowledge is still limited about the role of selection in the local adaptation of populations to climate changes and associated range shifts (De Lafontaine et al., 2018; Luqman et al., 2023), except for a few pioneering studies (Luqman et al., 2023; The 1001 Genomes Consortium, 2016).

Whole-genome polymorphism data from large population samples can be powerful for studying local adaptation in response to climate change in various ways (Bamba et al., 2019; Bourgeois and Warren, 2021; Weigel and Nordborg, 2015). First, the advance of statistical modeling based on coalescent theory enabled us to infer the demographic history of populations, including historical changes in effective population size and the timing of population divergence during glacial cycles. Second, genome-wide association studies (GWAS) can identify genes underlying phenotypes or other characteristics, such as the climate of sampled locations (Hancock et al., 2011; Lee et al., 2017; Rellstab et al., 2015; Savolainen et al., 2013; The 1001 Genomes Consortium, 2016). Third, genome-wide selection scans can detect loci under various selection modes, including selective sweeps and divergent selection.

The Japanese archipelago is an ideal area to study the mechanisms of how populations adapt to local climatic changes and associated range shifts, as it stretches 3,000 km from northeast to southwest over a wide range of climatic zones from subarctic to subtropical and has a complex mountainous topography in Central Japan (Ohsawa and Ide, 2011). Population structure analyses of several widely distributed species in the Japanese archipelago suggested range shifts during glacial cycles (Iwasaki et al., 2012; Magota et al., 2021; Ohsawa and Ide, 2011; Sakaguchi et al., 2018), but investigations on the climate adaptation at a region-wide scale are still scarce. This study focused on *Arabidopsis halleri,* a perennial herb distributed in East Asia and Europe. This species has been used extensively for evolutionary and ecological studies (reviewed in Honjo and Kudoh, 2019; Koch, 2018), such as self-incompatibility (Castric and Vekemans, 2004; Durand et al., 2020), heavy metal hyperaccumulation (Krämer, 2010; Stein et al., 2017), and adaptation to high altitudes (Kubota et al., 2015; Yoshida et al., 2023; Yumoto et al., 2021). Five subspecies are recognized in *A. halleri*: *A. halleri* subsp. *halleri*, *A. halleri* subsp. *tatrica*, *A. halleri* subsp. *dacica*, *A. halleri* subsp. *ovirensis*, and *A. halleri* subsp. *gemmifera* (Honjo and Kudoh, 2019). *A. halleri* subsp. *gemmifera* is distributed throughout Japan, Korea, north-eastern China, and the Russian Far East, while the other four subspecies are distributed in Europe. In Japan, *A. halleri* subsp. *gemmifera* is observed on the four main islands (Kyushu, Honshu, Shikoku, and Hokkaido). Given its wide distribution in the diverse climates across the Japanese archipelago, *A. halleri* can serve as a model species to study how plant populations are locally adapted to diverse climates and their changes during glacial cycles. Population structure and demographic histories are the fundamental basis for studying local adaptation, but the information is currently limited except for a few studies, including the one based on microsatellite markers (Sato and Kudoh, 2014) or the one on a microgeographic scale (Kubota et al., 2015; Yoshida et al., 2023).

In this study, using the whole-genome re-sequencing data of 141 Japanese *A. halleri* individuals and an outgroup of 16 European individuals, we aimed to understand how *A. halleri* adapted to local climates during glacial cycles through a population genomics approach. Specifically, we addressed the following questions:

1) How are Japanese populations of *A. halleri* genetically structured, and what demographic processes, such as population size fluctuations and divergence, formed the current pattern of population structure?
2) Which loci are associated with bioclimatic variables of sampled locations, and do these loci tend to be under selection? If so, in which subpopulation selection may have acted?
3) How have the potential habitats of *A. halleri* shifted since the Last Glacial Period?

## Results

### Population structure of A. halleri in Japan

We re-sequenced the genomes of 141 individuals from *A. halleri* wild populations distributed across Kyushu, Honshu, and Hokkaido islands in Japan using short reads (Supplementary Table S1). As outgroups, we also obtained sequence data of twelve and four individuals from 10 Central European populations and four Romanian populations, respectively. We mapped short reads to the *A. halleri* v2.03 assembly (DOE-JGI, http://phytozome.jgi.doe.gov/), obtaining 1,506,831 SNPs in total after variant calling and filtering. Network analysis using this SNP set revealed a genetic structure in accordance with geography: A cluster of Japanese *A. halleri* was distinct from Central European and Romanian clusters, and among the Japanese populations, Northern Japan (NJ) individuals formed a single group, separated from other individuals of Western Japan (WJ), Kansai Region (KR), and Central Japan (CJ) subpopulations (Fig. 1A). A clustering analysis using the non-negative matrix factorization (sNMF) also revealed the population structure that reflects geography in Japan (Fig. 1B, Supplementary Fig. S1A). The clustering pattern in *K* = 5 was largely concordant with the network-based analysis (Fig. 1B), while the best-supported cluster number was *K* = 7 based on the cross-entropy criterion (Supplementary Fig. S1B). These results overall suggest that Japanese *A. halleri* are geographically differentiated across the archipelago. For the following analyses of demographic history, individuals were classified into five subpopulations based on a maximum factor under *K* = 5 (Fig. 1C): Europe (EU), Western Japan (WJ), Kansai Region (KR), Central Japan (CJ), and Northern Japan (NJ).

**Fig. 1.**
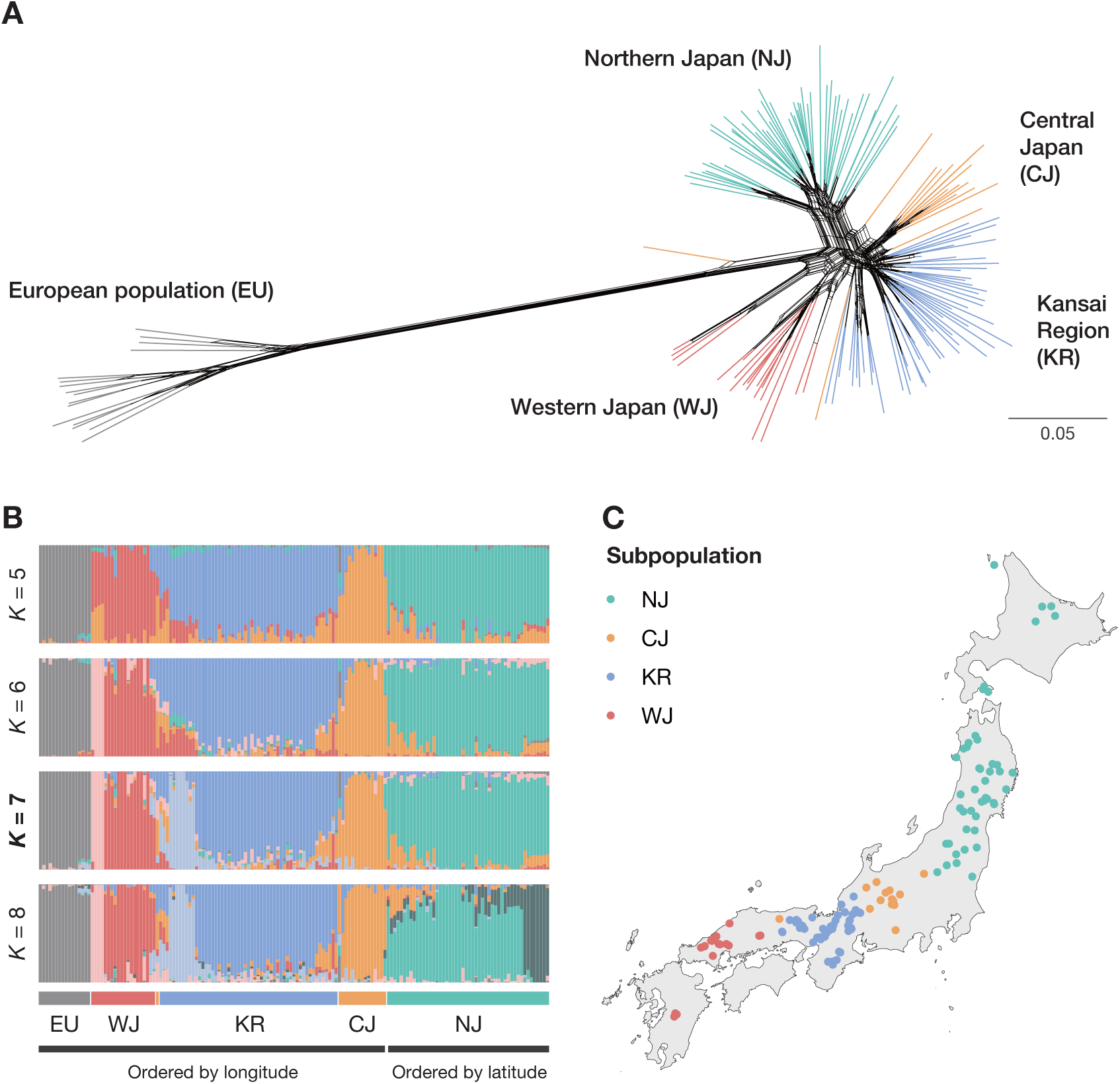
Population structure of *Arabidopsis halleri*. (A) A phylogenetic network generated by SplitsTree4 v.4.19.2 (Huson and Bryant, 2006). Scale bar indicates proportion of positions with different alleles. (B) Clustering by non-negative matrix factorization using the R package LEA v.3.10.2 (Frichot et al., 2014; Frichot and François, 2015). Results for *K* = 2–12 are available in Supplementary Fig. S1. Individuals are ordered by longitude for the EU, WJ, KR, and CJ subpopulations and by latitude for the NJ subpopulation. (C) Geographic distribution of Japanese individuals with the classification based on a maximum factor under *K* = 5.

### Demographic history and population subdivisions

We employed the approximate Bayesian computation (ABC) analysis to infer demographic history and divergence time among subpopulations. In this analysis, our primary focus was to obtain the approximate timescale of divergence time between subpopulations. Therefore, while more complex scenarios involving migrations and hybridizations could be considered, we examined four relatively simple scenarios consistent with the patterns of hierarchical clustering of population structure (Supplementary Fig. S2A; Fig. 1). Among them, scenario 3 was chosen as the best scenario with a posterior probability of 0.814, in which the WJ subpopulation first diverged from other subpopulations. Under scenario 3, the split time between the EU and other subpopulations (t4) was estimated to be 71,825.5 [95% credible interval of 35,370–140,944] generations ago (Fig. 2A, Supplementary Fig. S2, Supplementary Table S2). Assuming a generation time of two years (Honjo and Kudoh, 2019; Roux et al., 2011; Sato and Kudoh, 2014), this estimate corresponds to 143,651 [70,740 – 281,888] years ago. This time estimate of the Europe–Japan split within *A. halleri* was significantly smaller than the estimated divergence time between *A. halleri* and its closest relative, *A. lyrata,* which was about 337,000 [272,800–438,200] years ago (Roux et al., 2011). All the split events within Japanese subpopulations were inferred to have occurred during the Last Glacial Period (LGP; about 115,000–11,700 years ago) (EPICA community members, 2004): t3 = 52,322.2 [27,668–79,831.2], t2 = 35,398.6 [12,350.1–63,958], and t1 = 28,247.8 [7,404.54–57,254.4] years ago, assuming a generation time of two years. The ancestral population of all Japanese subpopulations was estimated to have the smallest effective population size (*N*_e_) (N_A3_ = 25,120.5 [4,022.72–66,230]).

**Fig. 2.**
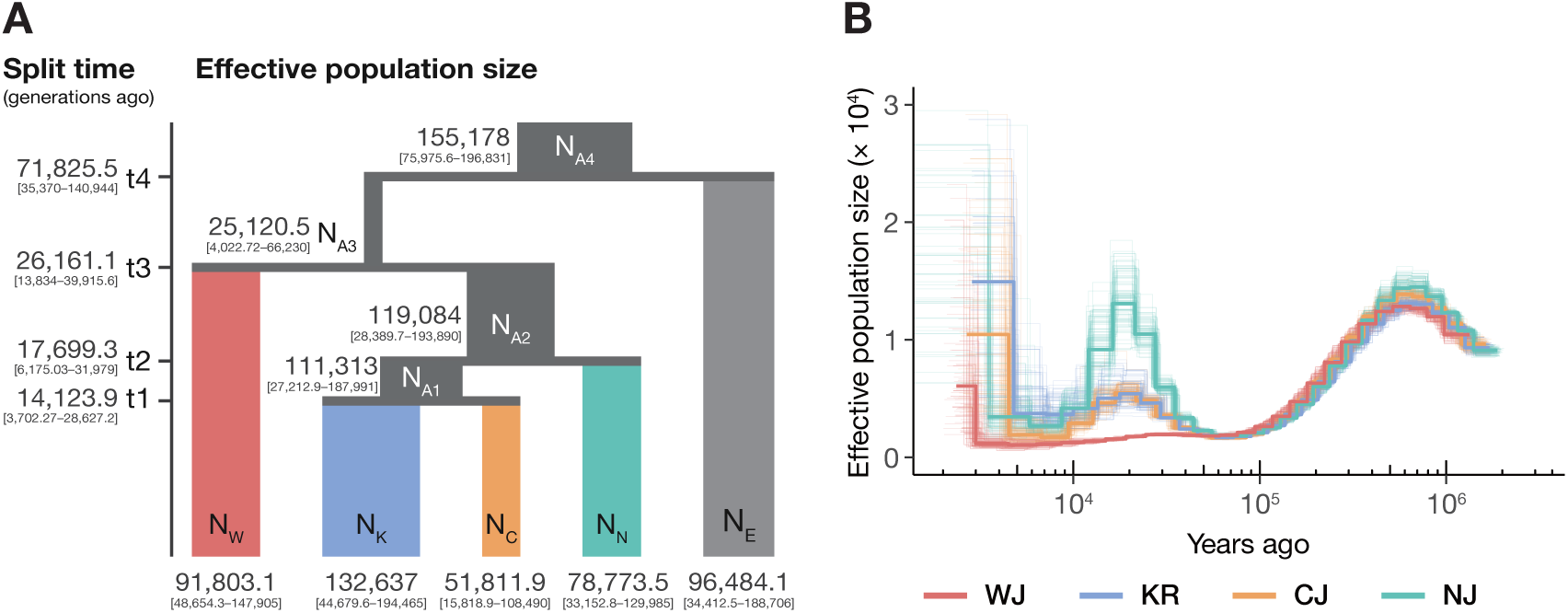
Demographic history of Japanese *Arabidopsis halleri*. (A) The best demographic model suggested by the approximate Bayesian computation analysis. The DIYABC-RF v.1.2.1 software was used (Collin et al., 2021). Effective population sizes of current and ancestral populations and split times are shown with 95% credible intervals. (B) The pairwise sequential Markovian coalescent analysis of four Japanese subpopulations (Li & Durbin 2011).

Although scenario 3 was the best model, scenario 2 was comparable to scenario 3 in terms of the classification vote, the number of times a scenario is selected in 1,000 random forest trees (Supplementary Fig. S2B). Nevertheless, both scenarios provided similar split times and effective population sizes, and all the split events within Japanese subpopulations were inferred to have occurred during the LGP under scenario 2 as well. (Table S2).

We also performed the pairwise sequential Markovian coalescent (PSMC) analysis to gain further insights into the demographic history within Japanese populations (Li and Durbin, 2011). We chose representative individuals for each Japanese subpopulation based on the result of clustering analysis to infer the historical changes in *N*_e_ (see Materials & Methods for details). Assuming a generation time of two years, all four subpopulations showed similar dynamics of *N*_e_ before ∼50 kya, but the more recent patterns were different between subpopulations: We observed the increase of *N*_e_ between 50–20 kya and the successive decrease until 7 kya in CJ, KR, and NJ subpopulations, while *N*_e_ of the WJ subpopulation declined for the period between 30–3 kya with a slight expansion before 30 kya (Fig. 2B). While we note that the PSMC-based estimate of recent past is relatively less reliable due to the limited number of independent coalescent events, different population size dynamics after 50 kya suggest the genetic differentiation between those subpopulations, which is consistent with the results of ABC analysis.

### Genomic signatures of climate adaptation in Japan

Next, we investigated the signatures of climate adaptation in the genomes of Japanese *A. halleri*. Our approach is summarized in Fig. 3A. First, we identified loci associated with local climate through genome-wide association mapping. Second, we performed genome-wide selection scans based on the cross-population extended haplotype homozygosity (XP-EHH) to detect subpopulation-specific selection (Gautier et al., 2017; Gautier and Vitalis, 2012; Klassmann and Gautier, 2022; Sabeti et al., 2007). Then, we asked if climate-associated loci are enriched in the extreme tails of selection scans. The statistical significance of the enrichments was assessed by permutation tests, with a null hypothesis that the fraction of the extreme tails of XP-EHH scores in climate-associated loci is the same as the fraction in the whole genome (climate-associated and climate-non-associated loci) (see Materials & Methods for details). Thus, high enrichment scores indicate that climate-associate loci tend to overlap with the positively selected regions than expected by chance. Since the XP-EHH-based selection scan detects the direction of selection, we performed the enrichment analyses for all the pairwise comparisons of four subpopulations in Japan to detect climate-associated subpopulation-specific local adaptation.

**Fig. 3.**
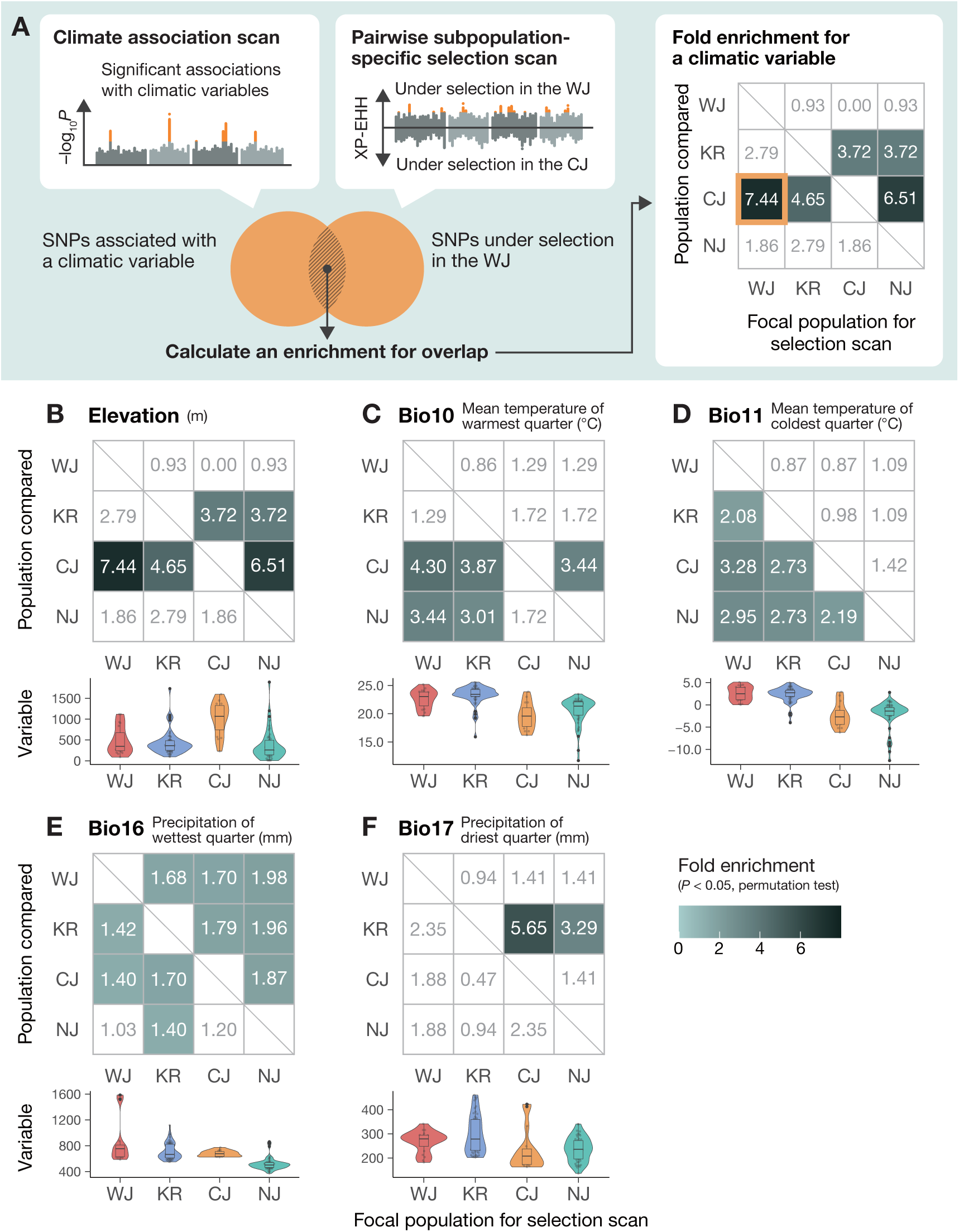
Selection enrichment analysis for climate adaptation. (A) A scheme of the analysis. A genome-wide association mapping was performed for each bioclimatic variable to find associated SNPs using the R package LEA v.3.10.2 (Gain and François, 2021). We also performed genome-wide selection scans based on the cross-population extended haplotype homozygosity (XP-EHH) for all pairwise combinations of subpopulations using the R package rehh v.3.2.2 (Gautier et al., 2017). Positive and negative values of XP-EHH indicate the direction of selection, providing information about which subpopulation selection may have acted. Then, fold enrichments of the overlap were calculated with their statistical significance based on permutation tests. High enrichment scores indicate that climate-associate loci tend to overlap with the positively selected regions than expected by chance. Enrichments and distributions of variables between subpopulations for elevation (B), bio10 (mean temperature of warmest quarter [°C]) (C), bio11 (mean temperature of coldest quarter [°C]) (D), bio16 (precipitation of wettest quarter [mm]) (E), and bio17 (precipitation of driest quarter [mm]) (F). (B-F) Because XP-EHH scores are calculated for all pairwise combinations of subpopulations, enrichments are also obtained for each focal subpopulation for selection scan (horizontal axis) and each subpopulation in comparison (vertical axis).

To perform these selection analyses, information on the population structure is required.

Because these analyses focus on Japanese populations, we re-analyzed population structure only with Japanese individuals (Supplementary Fig. S3A). We confirmed that the pattern of clustering was fairly close to the one when including European populations, and the individuals were classified into four Japanese subpopulations based on the maximum factor under *K* = 4: WJ, KR, CJ, and NJ (Supplementary Fig. S3B). Taking the information of population structure into account, we performed GWAS for elevation, 19 bioclimatic variables, and seven principal components of bioclimatic variables (PC1–PC7) using latent factor mixed models (LFMM) (Caye et al., 2019; Gain and François, 2021). These environmental variables were highly variable between four subpopulations (Fig. 3, Supplementary Fig. S4). The number of peaks satisfying genome-wide significance (4-kb windows containing SNPs with *P* < 0.05 after Benjamini–Hochberg false-discovery rate correction) ranged from 0 to 2,518 among the variables (Supplementary Fig. S5, S6). Gene ontology (GO) enrichment analysis for those peaks showed the enrichment of several terms, including those related to ion transport and metabolic processes (Supplementary Fig. S7).

To detect subpopulation-specific selection, we performed genome-wide selection scans based on the XP-EHH for all the pairwise comparisons of four subpopulations in Japan (Gautier et al., 2017; Gautier and Vitalis, 2012; Klassmann and Gautier, 2022; Sabeti et al., 2007). We first calculated the integrated extended haplotype homozygosity of a single nucleotide polymorphism site (iES), representing the average length of shared haplotypes (Tang et al., 2007). We then calculated cross-population extended haplotype homozygosity (XP-EHH) based on the cross-comparison of two populations to discover alleles with near-fixation in one population (Sabeti et al., 2007). XP-EHH values were calculated for each SNP with pairwise comparisons between the four subpopulations (Supplementary Fig. S8). Then, we performed enrichment analyses to test whether environmentally associated peaks are enriched in regions under subpopulation-specific selective sweeps. We found significant enrichments of 4-kb windows containing SNPs associated with elevation and 14 bioclimatic variables in the top or bottom 2.5% tails of the XP-EHH scores in at least one subpopulation comparison (Fig. 3, Supplementary Fig. S9, S10). As an alternative criterion for population differentiation, we also calculated *F*_ST_ between four subpopulations, detecting significant enrichments of windows containing SNPs associated with elevation and 15 bioclimatic variables in the top 5% of *F*_ST_ tails (Supplementary Fig. S11, S12, S13). Elevation-associated SNPs showed strong enrichments of the XP-EHH scores when focusing on WJ, KR, NJ subpopulations in comparison with the CJ subpopulation, which had the highest elevation level (Fig. 3B). SNPs associated with bio10 (mean temperature of warmest quarter) were also enriched in the XP-EHH scores when focusing on WJ and KR subpopulations in comparison with CJ and NJ subpopulations that had lower values of bio10 (Fig. 3C). Similarly, the association with bio11 (mean temperature of coldest quarter) showed significant enrichments in the XP-EHH scores when more southern subpopulations were compared with northern ones (Fig. 3D). In contrast, SNPs associated with bio16 (precipitation of wettest quarter) and bio17 (precipitation of driest quarter) tended to be enriched when focusing on more northern subpopulations compared with southern ones (Fig. 3E, F). These enrichment analyses demonstrate the prevailing selection signatures for climate-associated loci, particularly selection toward a warmer climate in southern subpopulations (WJ and KR) and a drier environment in northern subpopulations (CJ and NJ).

Here, we highlight a few genes showing associations with bioclimatic-associated SNPs and signatures of selection, particularly focusing on those that showed enrichments in more than two comparisons of subpopulations (Fig. 4A, Supplementary Table S3). Among the peaks associated with bio10 (mean temperature of the warmest quarter), two windows on chromosome 7 were commonly included in the 2.5% tail of XP-EHH scores in WJ, KR, and NJ subpopulations when compared with CJ populations (Fig. 4A). The SNP with the strongest association was located in the downstream of the gene Ah7G32100, annotated as a homolog of *A. thaliana HEAT INDUCIBLE LIPASE1*, and showed a clear geographic cline in the allele distribution (Fig. 4B). We also found three peaks associated with bio17 (precipitation of driest quarter) and under selection in the CJ and NJ subpopulations in comparison to the KR subpopulation (Fig. 4A). The SNP with the strongest association on chromosome 8 was an upstream gene variant of Ah8G12760, which also showed a geographic cline in the allele distribution (Fig. 4B).

**Fig. 4.**
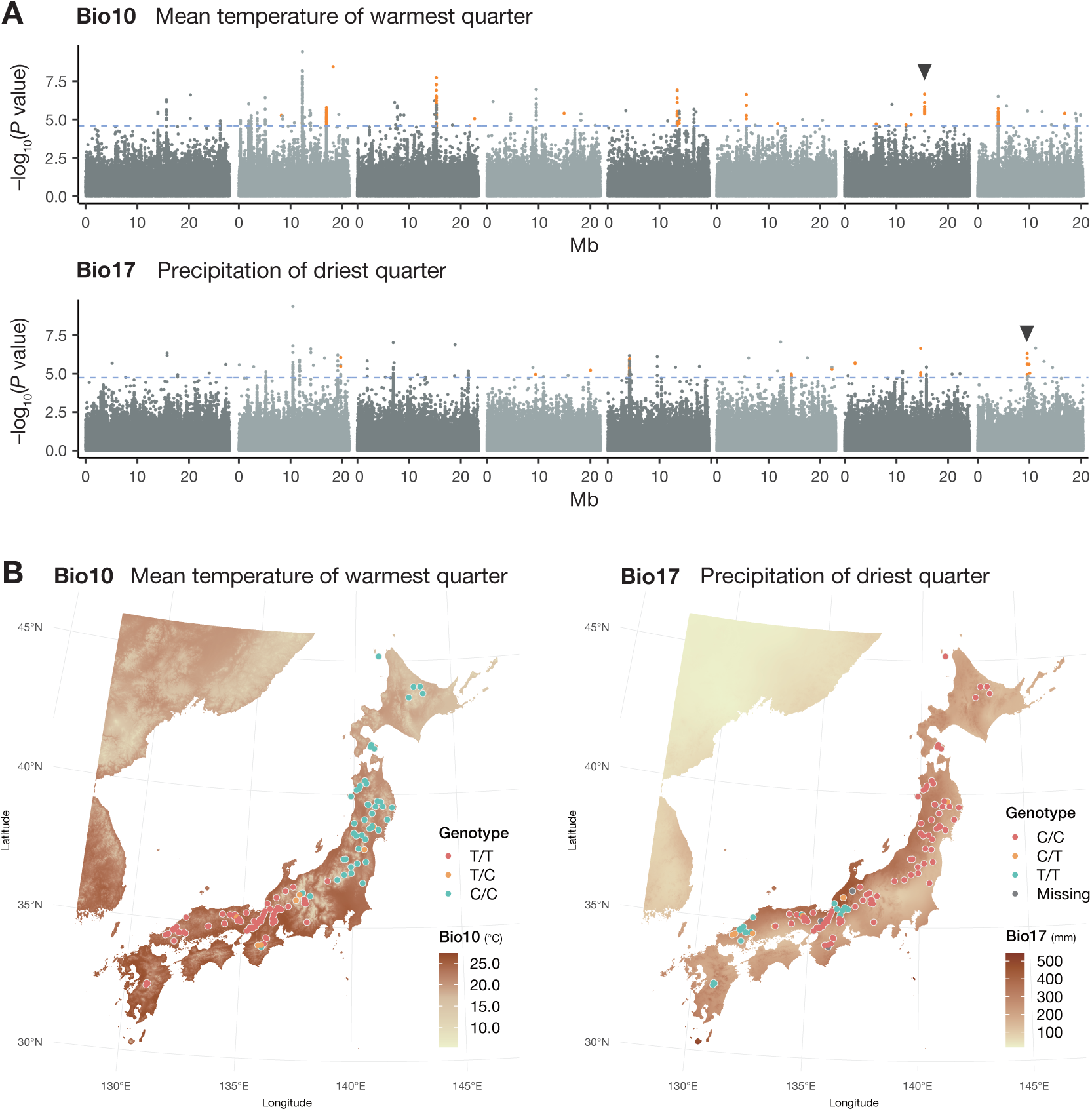
Climate-associated loci and their geographic distributions. (A) Manhattan plots for bio10 (mean temperature of warmest quarter) and bio17 (precipitation of driest quarter). Horizontal dashed lines indicate *P* = 0.05 after Benjamini–Hochberg false-discovery rate correction. Orange dots indicate SNPs associated significantly with bioclimatic variables and included in the 4-kb windows of selection scans that were significant in more than two combinations of subpopulations. SNPs indicated by arrowheads were used for plots of (B). SNPs on the assembled chromosomes are shown. (B) Geographic distributions of the alleles strongly associated with bio10 and bio17.

This gene was annotated as a homolog of *A. thaliana RAF10*, suggested to be involved in abscisic acid (ABA) response (Lee et al., 2015). In addition, we found a peak associated with bio11 (mean temperature of coldest quarter), which was under selection in the KR and CJ subpopulations compared with the NJ subpopulation. The most associated SNP was an upstream gene variant of Ah1G24310, a homolog of *EMBRYO DEFECTIVE 1507* in *A. thaliana*. Its mutant was reported to show reduced expression levels of *FLOWERING LOCUS C* (*FLC*) and early flowering (Mahrez et al., 2016). These genes are promising for climatic adaptation, particularly for warmer temperatures and lower precipitation.

### Historical changes in the suitable distribution area

As selection enrichment analysis suggested climate-driven selection, we next investigated the post-glacial changes in the distribution of Japanese *A. halleri* with a framework of ecological niche modeling (also called species distribution modeling). Ecological niche modeling is a numerical tool that combines observations of species occurrence with environmental estimates intended to predict distributions across landscapes, including extrapolation in space and time (Elith and Leathwick, 2009). Here, we first obtained distribution probabilities under the current climate conditions based on the sampling locations of Japanese *A. halleri* and six principal components of bioclimatic variables (see Materials & Methods for details). Then, we also developed distribution models under the Last Glacial Maximum (LGM; about 22,000 years ago) and Mid-Holocene (about 6,000 years ago). We found that potential habitats were predicted mainly in the lowlands of southern coastal regions in the LGM (Fig. 5A). In contrast, predictions based on Mid-Holocene and current climate conditions showed that potential habitats were shifted toward northern coastal and mountainous areas (Fig. 5B, C).

**Fig. 5.**
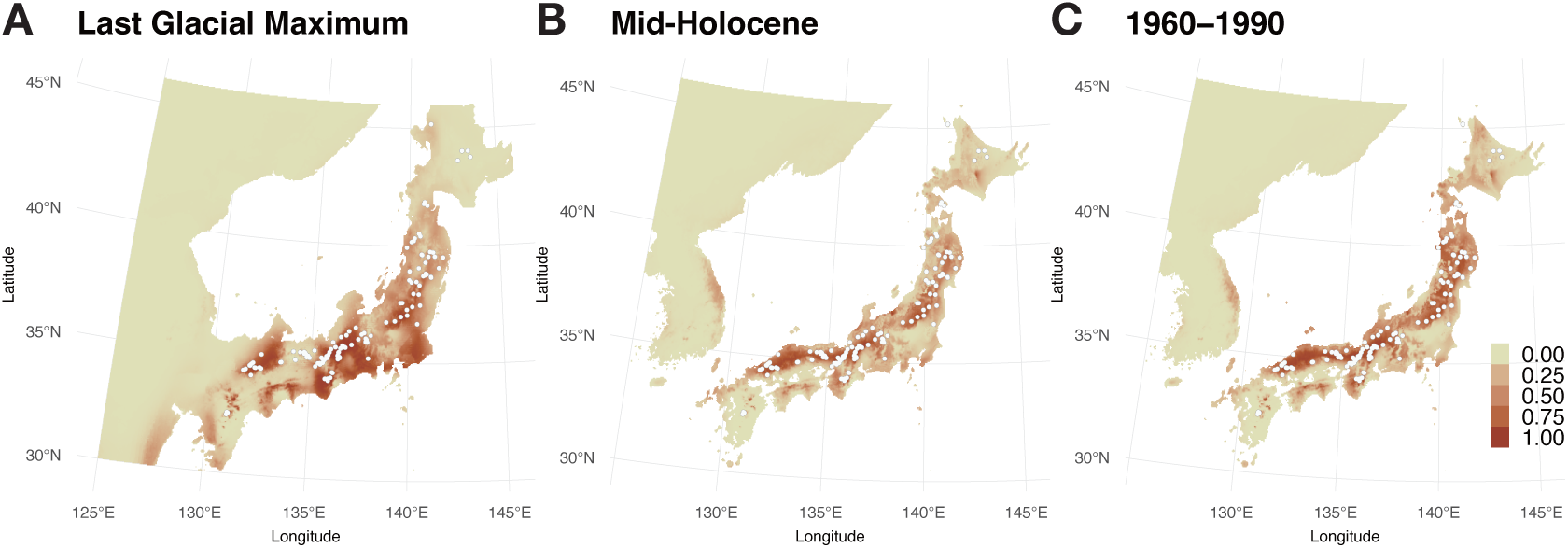
Ecological niche modeling of the potential habitat of *A. halleri*. MaxEnt v.3.4.4 was used (Phillips et al., 2024, 2004). Sample locations are indicated by open circles. The occurrence probabilities under the Last Glacial Maximum (about 22,000 years ago) (A), Mid-Holocene (about 6,000 years ago) (B), and the current (1960–1990) climate conditions (C) are shown.

## Discussion

### Genomic signatures of climatic adaptation during glacial cycles

In this study, we investigated population structure, demographic history, and signatures of climate-driven selection by exploiting genome-wide polymorphism data of 141 Japanese *A. halleri* individuals and European ones as outgroups. The ABC and the PSMC analyses consistently suggested the genetic differentiation between Japanese subpopulations since the LGP (Fig. 2), which would contribute to shaping the current pattern of population structure. Clonal reproduction and a lack of specific structure for long-distance dispersal reported in this species may also have facilitated the formation and maintenance of population structure (Honjo and Kudoh, 2019; Sato and Kudoh, 2014). Population demographic analysis revealed the population size fluctuations in the LGP, which was particularly prominent since the subpopulations started to diverge (∼50 kya). Together with the prediction by the ecological niche modeling that the potential habitat may have shifted from southern coastal regions to northern coastal and mountainous areas (Fig. 5), our analyses overall suggest that subpopulations in Japanese *A. halleri* have experienced population subdivisions, fluctuations, and distribution range shifts accompanied by the climatic change during and after the LGP, as also reported in other plants and animals (e.g., Comes and Kadereit, 1998; Hewitt, 2004; Qiu et al., 2011).

The combination of GWAS for bioclimatic variables and XP-EHH-based selection scans enabled us to detect signatures of subpopulation-specific selection with explicit climatic contexts (Fig. 3, Supplementary Fig. S9, S10). An advantage of using XP-EHH is that we can identify not only the adaptive divergence between subpopulations but also the direction of selection, *i.e.*, in which subpopulations selection might have acted. For example, temperature-associated GWAS peaks were enriched in the XP-EHH tail when focusing on southern subpopulations (WJ and KR), suggesting adaptation toward higher temperatures in those subpopulations (Fig. 3C, D). Similarly, the enrichment of GWAS peaks associated with the precipitation of the driest quarter was observed in the XP-EHH scores when focusing on Central Japan (CJ) and Northern Japan (NJ) populations, suggesting the adaptation toward the lower precipitation (Fig. 3F). Although our study aimed at detecting polygenic selection for local climates rather than screening genes, it is worth mentioning that several promising genes appeared to be linked to the associated SNPs, such as homologs of *HEAT INDUCIBLE LIPASE1* and *RAF10*, involved in ABA response, associated with the mean temperature of the warmest quarter and precipitation of the driest quarter, respectively. Given the subpopulation divergence since the LGP, such subpopulation-specific climatic adaptation for higher temperature or lower precipitation would have occurred during or after the LGP, possibly accompanied by population contraction and expansions. In *A. thaliana*, which experienced an expansion of distribution after the LGP, a study using more than 1,000 natural strains also identified SNPs associated with precipitation-related bioclimatic variables, and they showed a significant geographic gradient (Lee et al., 2017; The 1001 Genomes Consortium, 2016).

Except for those pioneering studies, attempts to identify loci potentially involved in adaptation are still scarce. Our study highlights the importance of integrating climate associations, selection scans, and population demographic analyses to identify genomic signatures of population-specific adaptation when comparing several genetic clusters. Such combined approaches would also be important for reducing false positives of environmental association analysis that are suggested to be high (Rellstab et al., 2015).

### Population structure and demography history of A. halleri

While there have been a few studies on population structure in Japanese *A. halleri*, they relied on a limited number of loci and sampling locations (Sato and Kudoh, 2014) or focused on a microgeographic scale (Kubota et al., 2015; Yoshida et al., 2023). In this study, by using full-genome re-sequencing data from region-wide samples, we revealed a population structure that showed a clear geographic pattern (Fig. 1). Based on the data of population structure, European *A. halleri* were inferred to have been distributed in multiple refugia in the LGP (Šrámková et al., 2019; Šrámková-Fuxová et al., 2017). Our population demography analysis suggests population subdivision during the LGP, consistent with the persistence of multiple refugia. Our ecological niche modeling also identified multiple potential habitats along the southern coastal regions during the LGM, which might have served as glacial refugia. Species occurring in similar geographic regions sometimes share similar patterns of population structure, which may reflect shared history during glacial cycles. The pattern of geographical differentiation of Japanese *A. halleri* appeared to be at least partially comparable with those of other plants widely distributed in Japan. The genetic boundary between northern and southern clusters similar to that of CJ and NJ subpopulations are identified in the phylogeographic analysis of several plant species in Japan (Iwasaki et al., 2012; Magota et al., 2021; Ohsawa and Ide, 2011). The phylogeographic analysis of the goldenrod *Solidago virgaurea* complex has identified similar genetic clusters across the Japanese archipelago (Sakaguchi et al., 2018). These shared population structures would imply the similarity of distribution shifts since the LGP and the survival in the same refugia (Iwasaki et al., 2012). The smallest *N*_e_ of ancestral population of all Japanese subpopulations in *A. halleri* may suggest a demographic bottleneck upon colonization into the Japanese archipelago, consistent with the migration to the archipelago through land bridges that has been suggested in some species (Ohsawa and Ide, 2011; Sakaguchi et al., 2018).

### Conclusion and future perspectives

In this study, the integrated analyses of population demography, climate associations, selection scans, and ecological niche modeling revealed the signatures of subpopulation-specific climatic adaptation, particularly for higher temperature or lower precipitation, which would have taken place during or after LGP, possibly accompanied by population contraction and expansions in Japanese *A. halleri.* Knowledge of the adaptation to past and current environments should also help us predict the evolutionary responses to future climate changes. Recent population genomic studies focusing on drought tolerance in *A. thaliana* predicted its distribution under future climate models based on the geographic information of adaptive alleles and field experiments (Exposito-Alonso et al., 2019, 2017). While there is a growing interest in using population genomics to predict future evolutionary responses, the importance of validating genomic predictions has been suggested (Capblancq et al., 2020). The functional relevance of several genes that appeared in our climate GWAS needs to be validated in *A. halleri*, and common garden experiments in multiple locations in Japan would be helpful for such validations. We would also like to note that not only adaptation to temperature or precipitation, the evolution of mating systems and polyploidy is often suggested to be associated with population demographic changes during LGP and post-glacial expansions (Durvasula et al., 2017; Novikova et al., 2017; Tsuchimatsu et al., 2012, 2010; Willi et al., 2022). Investigation of further phenotypic variation and its genetic basis will be valuable for a better understanding of climate adaptation mechanisms.

## Materials and Methods

### Sampling and whole-genome resequencing

In this study, we used whole-genome re-sequencing data of 141 Japanese and 16 European individuals (Supplementary Table S1). The 141 locations of Japanese samples largely cover the distribution range of *A. halleri* in Japan, according to the Global Biodiversity Information Facility (https://www.gbif.org/). Among these individuals, eight were sequenced by Kubota et al., 2015. For other individuals, whole-genome re-sequencing was performed by the following methods, depending on samples.

For 90 Japanese individuals, we sampled about 30 mg dried biomass of leaves per individual from herbarium specimens or from samples collected in each location, and genomic DNA was extracted using the DNeasy Plant Mini Kit (Qiagen, Hilden, Germany) (Supplementary Table S1). DNA quality, quantity, and integrity were assessed with a Nanodrop spectrophotometer (ThermoFisher Scientific, Darmstadt, Germany). After the fragmentation by the NEBNext dsDNA Fragmentase kit (Ipswich, MA, USA), libraries were prepared using the TruSeq DNA LT Sample Prep Kit (Illumina, San Diego, USA) according to the manufacturer’s instructions, and whole-genome sequencing with 100-bp single-end reads were performed using an Illumina HiSeq 2000 instrument (Illumina, San Diego, USA). For herbarium samples, the fragmentation procedure was not performed.

For other 43 Japanese individuals, we sampled about 30 mg dried biomass of leaves per individual, and genomic DNA was extracted using a modified cetyltrimethylammonium bromide method (Milligan, 1992). DNA quality, quantity, and integrity were assessed with a Qubit 2.0 fluorometer (Thermo Fisher Scientific, Waltham, MA, USA) and a Nanodrop spectrophotometer (ThermoFisher Scientific, Darmstadt, Germany). Library preparations and 100-bp paired-end sequencing was performed using a BGISEQ-500 next-generation sequencing platform (BGI, Shenzhen, China).

To obtain whole genome re-sequencing data of 16 European individuals, living *A. halleri* plants collected in the field (Stein et al., 2017) were maintained in a greenhouse and propagated vegetatively. We sampled 50 to 70 mg fresh biomass of young leaves per individual in a 1.5 mL polypropylene tube and shock-froze the tissues in liquid nitrogen. Leaf tissues were homogenized using a single metal bead (5 mm stainless steel ball) per tube in a Retsch mixer mill (Type MM 300, Retsch, Haan, Germany) for 1 min at 30 Hz, using adapters pre-cooled to -80°C. The homogenate was used to isolate DNA using the NucleoMag^®^ Plant kit (Macherey Nagel, Düren, Germany) on an epMotion 5075 robot according to the manufacturers’ instructions (Eppendorf, Hamburg, Germany). DNA quality, quantity, and integrity were assessed on 0.8% (w/v) TAE-agarose gels, with a Nanodrop 2000 spectrophotometer (ThermoFisher Scientific, Darmstadt, Germany) and a Qubit 2.0 Fluorometer (Invitrogen, Darmstadt, Germany) using the dsDNA BR Assay Kit. Per sample, 1 µg (50 µl) genomic DNA was used as an input for producing libraries using the Illumina TruSeq DNA PCR-Free Library Prep according to the manufacturer’s instructions. Libraries passing QC were diluted, pooled and dispatched to Novogene Europe (Cambridge, United Kingdom) for whole-genome sequencing with 150-bp paired-end reads at a targeted minimum depth of 20x genome coverage using an Illumina NovaSeq 6000 instrument (Illumina, San Diego, USA).

### SNP identification

Whole-genome short reads were trimmed using Trimmomatic 0.39 with the options of LEADING:5, TRAILING:5, SLIDINGWINDOW:4:15, and MINLEN for 30% of the read length (Bolger et al., 2014). To be consistent throughout the analysis, pair-end reads were concatenated and treated as single-end reads. We then mapped read files with BWA-MEM 0.7.17-r1188 (Li, 2013) to *A. halleri* v2.03 assembly (DOE-JGI, http://phytozome.jgi.doe.gov/), then sorted and indexed the bam files with samtools v.1.17 (Danecek et al., 2021). The average depths were calculated for each bam file using the samtools depth command. We used bcftools v.1.17 mpileup and call pipeline for the variant calling (Danecek et al., 2021). SNPs with quality ≤ 20, minor allele frequency ≤ 0.05, depth ≤ 1,500 or depth ≥ 4,300, and a fraction of missing individuals > 0.1 were filtered out using bcftools. We also conducted SNP identifications without European individuals using the same pipeline and filtering options except for depth (SNPs with depth ≤ 1,200 or depth ≥ 3,600 were filtered out). This set was used for population structure analysis specific to Japanese individuals, climate association mapping, and selection scans.

### Population structure and network analysis

Population structure analysis was conducted using non-negative matrix factorization (sNMF) using the snmf function (*K* = 1–12; 50 repetitions for each *K*) implemented in the R package LEA v.3.10.2 (Frichot et al., 2014; Frichot and François, 2015). We employed the cross-entropy criterion to determine the most supported number of *K*. A population structure analysis without European individuals was also performed (*K* = 1–12; 10 repetitions for each *K*), and the individuals were classified into four Japanese subpopulations according to the maximum factor of *K* = 4. Selection scans in Japanese subpopulations were based on this classification.

To analyze the genetic relationship between individuals, we generated a nexus file from the vcf file using vcf2phylip (Ortiz, 2019), which was used as an input for SplitsTree4 v.4.19.2 (Huson and Bryant, 2006).

### Inference of demographic history and population subdivisions

We employed the ABC analysis using the DIYABC-RF v.1.2.1 software (Collin et al., 2021). For the input for ABC analysis, 20,000 SNP sites with no missing individuals were randomly extracted from the filtered SNPs and converted using the vcf2diyabc.py script (https://github.com/loire/vcf2DIYABC.py). Based on population structure and network analysis, we assumed an evolutionary scenario in which all the subpopulations diverged simultaneously and three other bifurcating scenarios (Fig. 2; Supplementary Fig. S2). In the latter three scenarios, KR and CJ subpopulations were assumed to form a single clade, as they were suggested to be closely related in the population structure analysis. Because our primary focus was to obtain the approximate timescale of divergence time between subpopulations, we examined these relatively simple scenarios without migrations and hybridizations. Considered scenarios were consistent with the patterns of hierarchical clustering of population structure (Fig. 1B). The prior distributions were set to 10–200,000 for all the population size parameters, 20,000–500,000 for the split time of the EU population, and 10–50,000 for other timings. After running 80,600 simulations, the best scenario was chosen based on the number of classification vote, and parameters were estimated for each scenario with 1,000 out-of-bag testing samples and 1,000 trees for random forest analysis.

Historical effective population sizes of four Japanese subpopulations were inferred using the PSMC model (Li and Durbin, 2011), implemented in psmc v.0.6.5-r67. We first selected representative individuals with the highest portion of the subpopulation-specific elements in the clustering results of sNMF with an average depth > 28, calculated by the SNP identification pipeline. We mapped reads of these individuals to hardmasked *Arabidopsis halleri* v2.03 assembly (DOE-JGI, http://phytozome.jgi.doe.gov/) using BWA-MEM and identified SNPs using bcftools mpileup and call pipeline. With the average depth calculated from these bam files using the samtools depth command, we converted them into a fastq file using vcfutils.pl with the option -d set to one-third and -D set to twice the average depth, which was then converted using fq2psmcfa with the -q20 option. We conducted the psmc analysis with 100 bootstrap runs using the options of -N25, -t15, -r5, and -p “4 + 25*2 + 4 + 6”. The results were plotted assuming a mutation rate of 7.1 × 10^–9^ per site per generation (Ossowski et al., 2010) and a generation time of two years.

### Genome-wide scans for climate associations

We conducted a genome-wide association mapping to identify loci associated with environmental gradients. We used elevation and bioclimatic variables of the places of origin based on the WorldClim v1.4 historical climate data (Hijmans et al., 2005) obtained using the raster package (Hijmans, 2024) in R. Details about bioclimatic variables are available in Supplementary Table S4. These variables were at the 30 arc-second resolution data. As these variables were correlated with each other, we performed principal component analysis (PCA) for 19 bioclimatic variables of the individuals using the prcomp function in R. We used seven principal components (PC1–PC7) for further analysis, which cumulatively explained 99.27% of the variance (Supplementary Fig. S14). We then imputed missing genotypes using the impute function in the R package LEA (Gain and François, 2021) based on the result of population structure analysis within Japan. Genome-wide association mapping was performed using LFMM implemented in the R package LEA lfmm2 function (Caye et al., 2019; Gain and François, 2021), specifying the number latent factor *K* = 7. We then identified 4-kb windows containing SNPs with *P* < 0.05 using Benjamini– Hochberg false-discovery rate (FDR) correction. This window size was based on the genome-wide pattern of linkage disequilibrium decay within 4 kb, estimated by PopLDdecay (Zhang et al., 2019) (Supplementary Fig. S15).

### GO enrichment analysis

We performed GO term enrichment analysis for the genes closely linked to climate-associated SNPs. We first identified genes overlapped with the 4-kb windows containing SNPs with *P* < 0.05 (FDR correction). The GO terms were obtained from *A. halleri* v.2.1.0 annotation data (DOE-JGI, http://phytozome.jgi.doe.gov/). GO term enrichment analysis for terms including more than nine genes was performed using topGO (Alexa et al., 2006) in R, with the weight01 algorithm and the significance level of *P* < 0.05 (Fisher’s exact test).

### Genome-wide selection scan

We performed genome-wide selection scans based on the cross-population extended haplotype homozygosity (XP-EHH) to detect subpopulation-specific selections. To calculate XP-EHH, we used the R package rehh v.3.2.2 (Gautier et al., 2017; Gautier and Vitalis, 2012; Klassmann and Gautier, 2022; Sabeti et al., 2007). Positive and negative values of XP-EHH reflect the directions of selection based on the difference in haplotype homozygosity between subpopulations. We first calculated iES statistics for SNPs on chromosome 1–8 using the scan_hh function in the pairwise combination of four Japanese subpopulations. We then calculated XP-EHH using the ies2xpehh function and extracted maximum and minimum scores in the 4-kb windows. We also calculated weighted *F*_ST_ in the 4-kb window using VCFtools v. 0.1.16 —weir-fas-pop option (Danecek et al., 2011).

We then calculated the enrichment of environmentally associated windows in the top or bottom extreme tail of the XP-EHH and *F*_ST_ values. The threshold for the XP-EHH tail was set to 2.5% for both sides. Windows containing climatic-associated sites with *P* < 0.05 (FDR correction) were considered. For the *F*_ST_ values, we set the threshold to the top 5%.

We conducted permutation tests to determine the statistical significance of fold enrichment.

For each of the permutations, we calculated fold enrichment between climate-associated window sets and randomly shifted XP-EHH window sets, preserving the relative position of the windows to take linkage disequilibrium between loci into account (Atwell et al., 2010; Horton et al., 2012; Nordborg et al., 2005; Tsuchimatsu et al., 2020). We conducted 1,000 times of permutation for each combination of bioclimatic variables and populations.

We also analyzed genes overlapping with the significant windows in climatic associations and selection scans. We especially focused on the significant windows in more than two pairwise combinations of subpopulations. The gene annotation was based on *Arabidopsis halleri* v.2.1.0 annotation data (DOE-JGI, http://phytozome.jgi.doe.gov/), and information on genes linked to each SNP was obtained by SnpEff v.5.1d (Cingolani et al., 2012). We plotted the geographic distribution of alleles on the loci of interest using the vcfR package (Knaus and Grünwald, 2017), with bioclimatic variables converted using the raster package (Hijmans, 2024) in R.

### Ecological niche modeling

We conducted ecological niche modeling using MaxEnt v.3.4.4 (Phillips et al., 2024, 2004) based on the 2.5 arc-minute resolution bioclimatic variables obtained from WorldClim v1.4 (Hijmans et al., 2005). These variables were cropped into the geographical range of 125–146°E, 30–46°N using a raster package (Hijmans, 2024) in R. As the bioclimatic variables are correlated with each other, we first conducted PCA for historical bioclimatic data using prcomp function in R to prevent overfitting. We then used PC1–PC6, which cumulatively explained 99.08% of variance as environmental layers (Supplementary Fig. S16). These components were projected to paleo climate variables at Mid-Holocene (about 6,000 years ago) and LGM (about 22,000 years ago) on WorldClim v1.4, which were then used as projection layers. Analysis was performed based on sample locations with default parameters.

### Data Availability

The sequence data underlying this article are available in European Nucleotide Archive (accession number: PRJEB72840).

## Acknowledgments

Authors thank Yu Okamura for helpful discussions, Naoko Ishikawa, Takaya Iwasaki, Masayuki Maki, Shuichi Nemoto, Shota Sakaguchi, Mahoro Suzuki, Akitomo Uchida, Yoichi Watanabe, and Yoshiharu Yamamoto for collecting samples, Atsushi Ebihara, Noriaki Murakami, Jin Murata, Hidetoshi Nagamasu, Hiroyoshi Ohashi, Takashi Shiga, and Hideki Takahashi for providing herbarium specimens, and Motomi Ito, Yutaka Suzuki, and Sumio Sugano for supporting genome sequencing.

## Author Contributions

RAS, SK, S-I M, and TT conceived and designed the study. SK, VK, UK, and S-I M generated the sequence data. RAS analyzed the data with help from VC and TT. RAS and TT wrote the paper with inputs from all authors.

