## Supplementary Figures 1-16 for "Population genomics reveals demographic history and climate adaptation in Japanese *Arabidopsis halleri*"

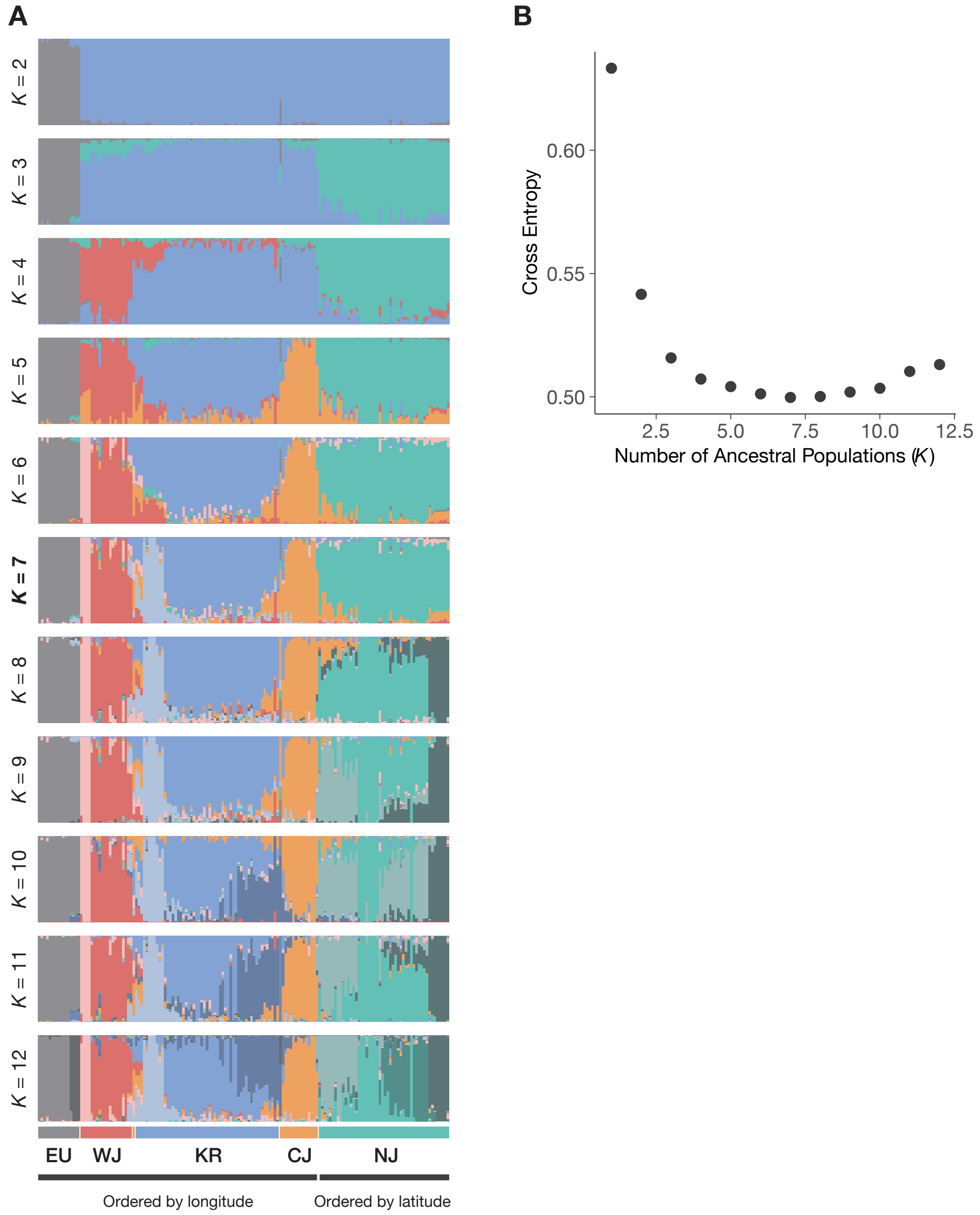


Supplementary Fig. S1. Clustering by non-negative matrix factorization using the R package LEA v.3.10.2 (Frichot et al., 2014; Frichot and François, 2015). (A) Results for $K$ = 2–12. Individuals are ordered by longitude for EU, WJ, KR and CJ subpopulations and by latitude for the NJ subpopulation. (B) The cross entropy for each $K$.

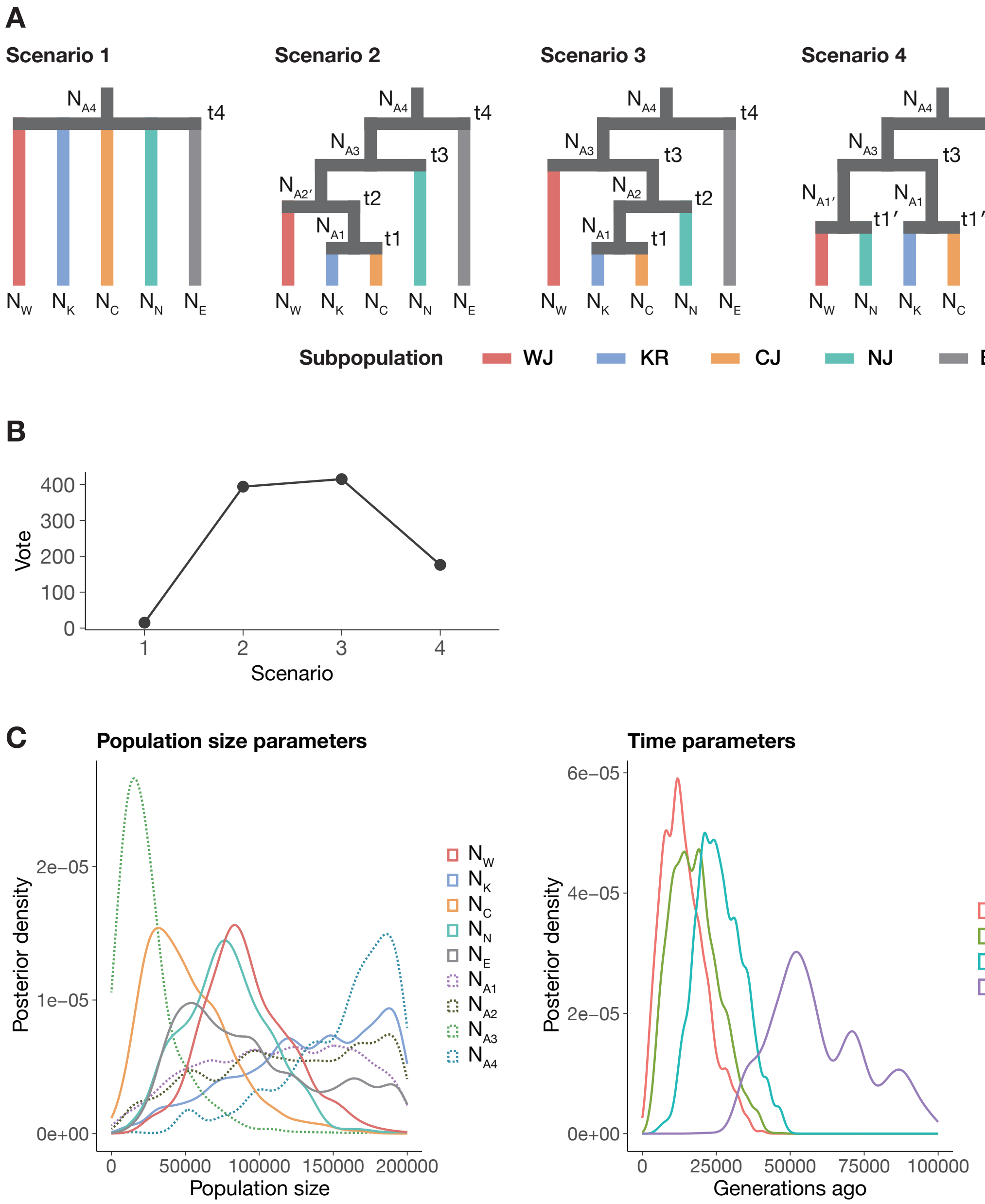


Supplementary Fig. S2. The Approximate Bayesian Computation analysis using the DIYABC-RF v.1.2.1 software (Collin et al., 2021). (A) Four scenarios considered in this study, which are consistent with the patterns of hierarchical clustering of population structure. Characters starting with "N" indicate the effective population size and are placed at the bottom of the current populations and the left of the respective ancestral populations. Characters starting with "t" indicate split time and are positioned to the right above the events. Inferred value for each parameter is shown in Supplementary Table S2. (B) Number of classification vote for each scenario, which represents the number of times that the scenario was selected in 1,000 random forest trees. Scenario 3 was chosen as the best scenario with the highest number of votes. (C) Inferred parameter distributions under the scenario 3.

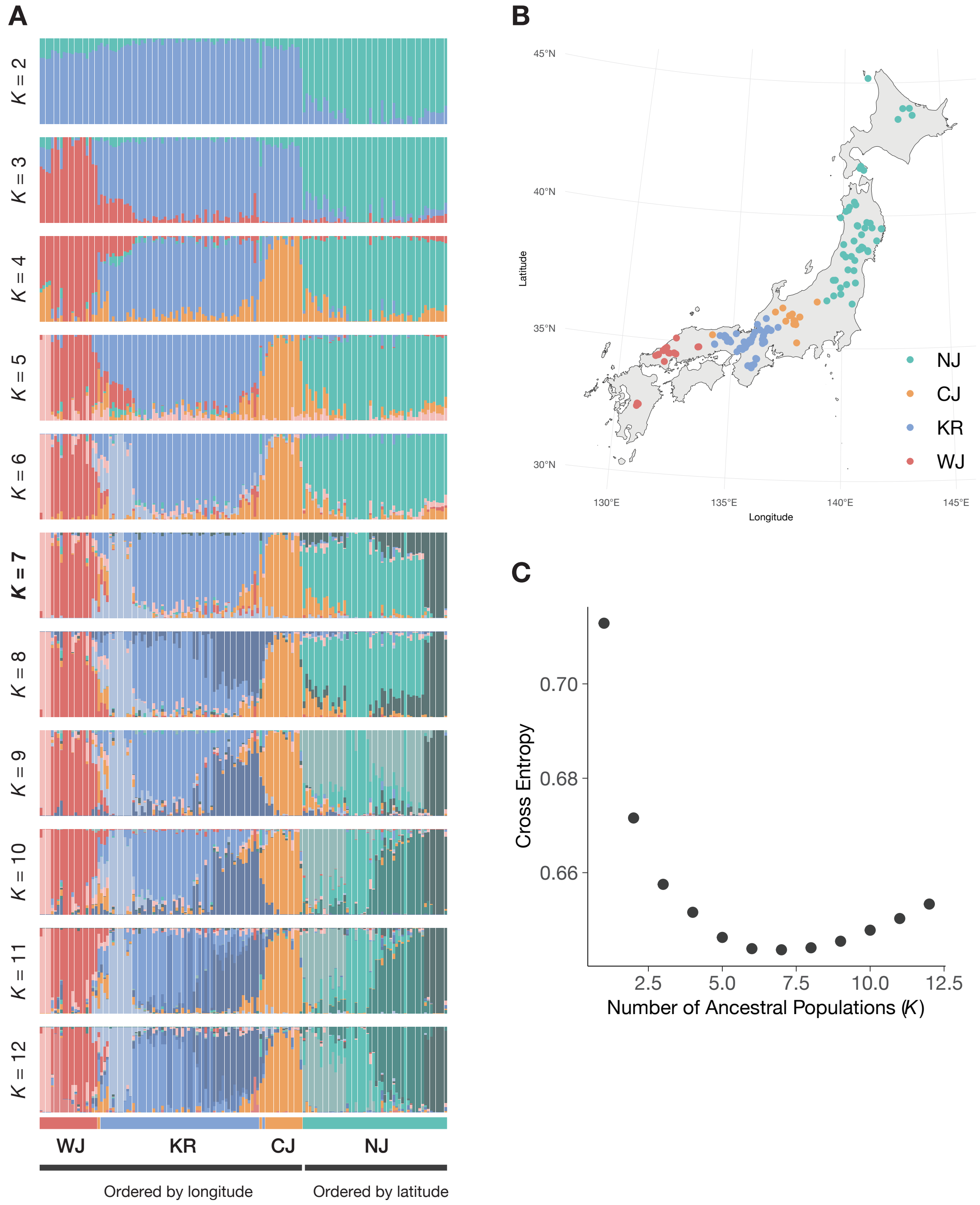


Supplementary Fig. S3. Clustering by non-negative matrix factorization using the R package LEA v.3.10.2 (Frichot et al., 2014; Frichot and François, 2015) only with Japanese individuals. (A) Results for $K = 2–12$. Individuals are ordered by longitude for WJ, KR and CJ subpopulations and by latitude for the NJ subpopulation. (B) Geographic distribution of Japanese individuals with the classification based on a maximum factor under $K = 4$. (C) The cross entropy for each $K$.

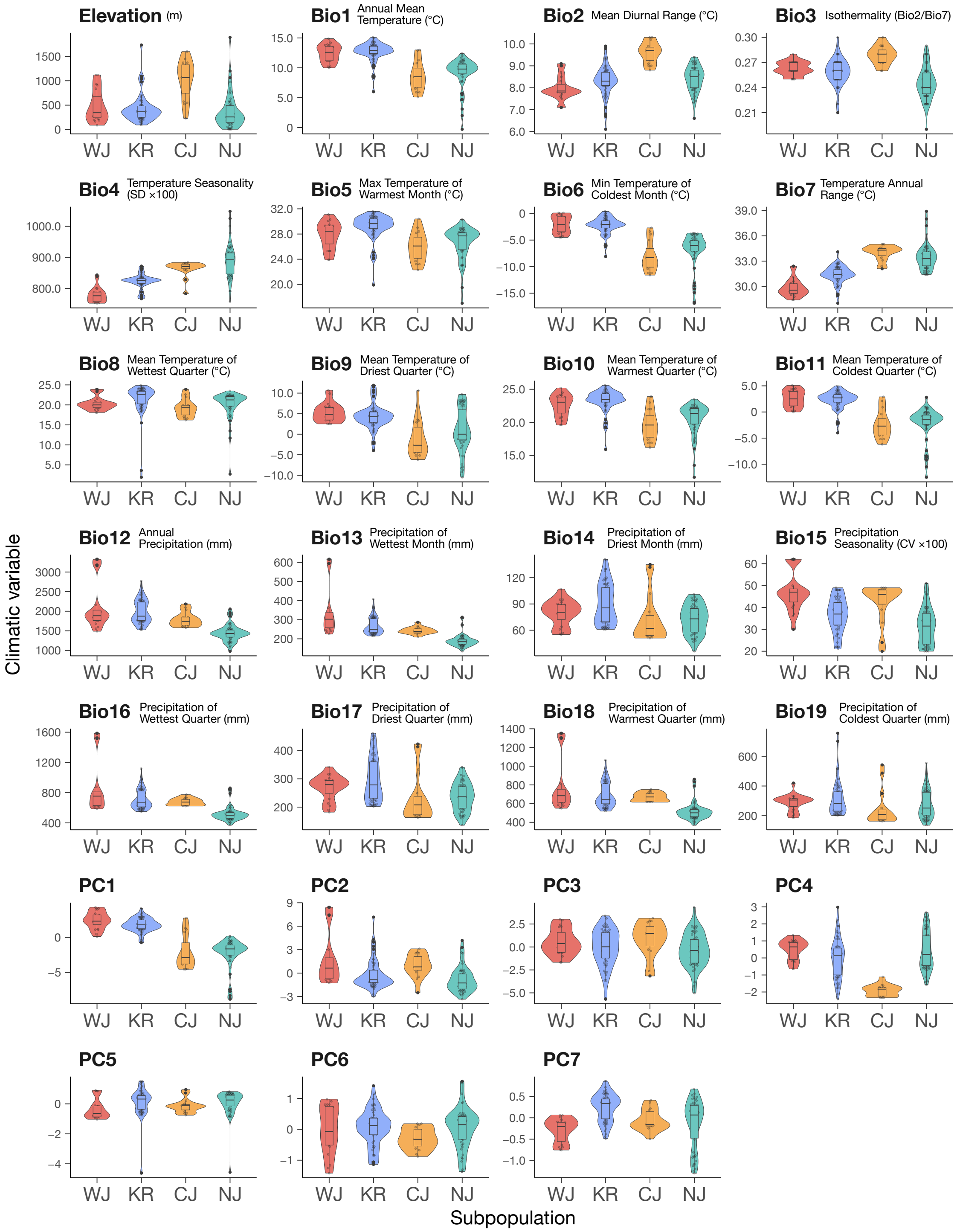


Supplementary Fig. S4. Distributions of bioclimatic variables and their principal components for each subpopulation. See Supplementary Table S4 for the list of bioclimatic variables.

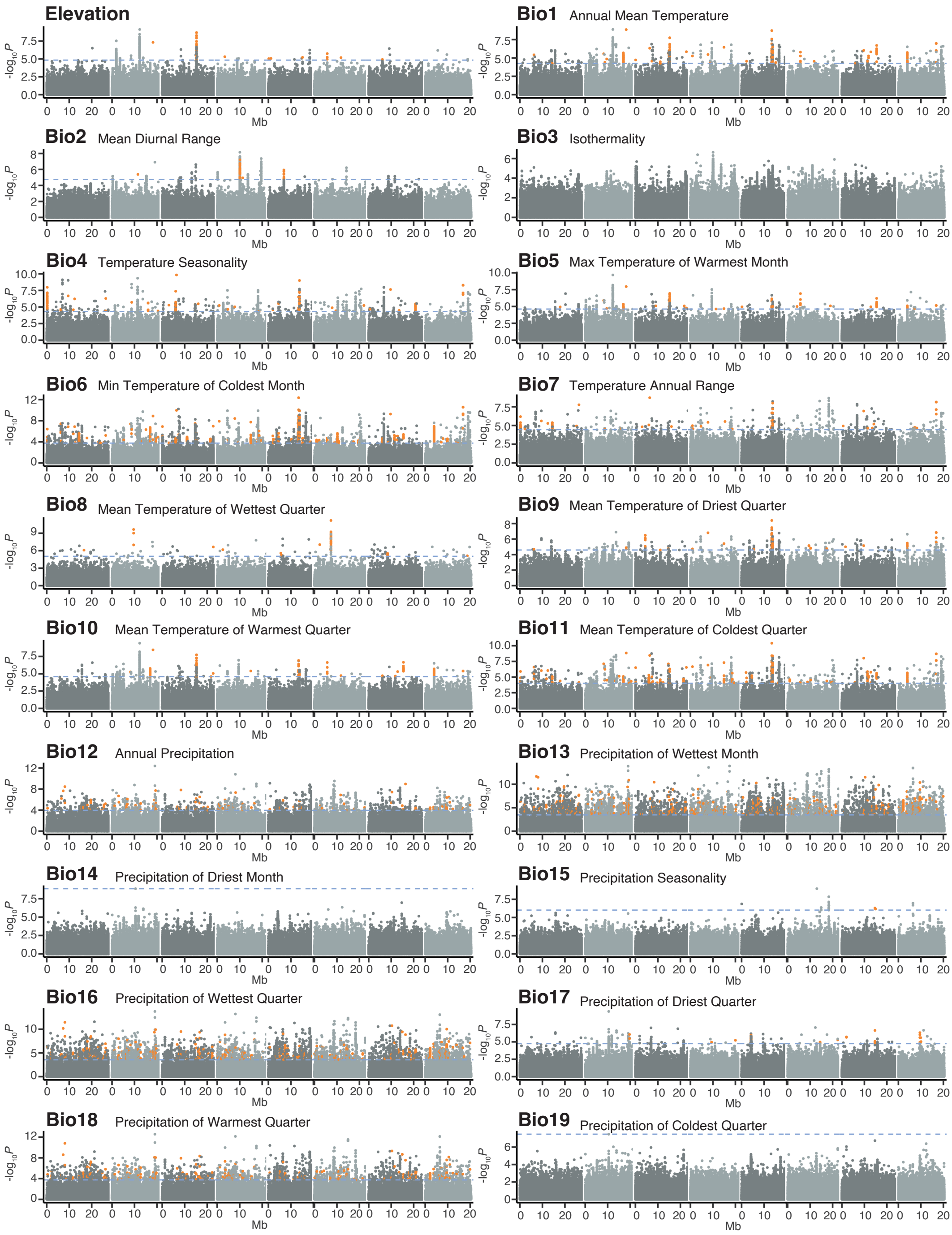


Supplementary Fig. S5. Manhattan plots for all the bioclimatic variables. Horizontal dashed lines indicate $P = 0.05$ after Benjamini–Hochberg false-discovery rate correction. Orange dots indicate SNPs associated significantly with bioclimatic variables and included in the 4-kb windows of selection scans that were significant in more than two combinations of subpopulations. See Supplementary Table S4 for the list of bioclimatic variables. Latent factor mixed models in the R package LEA was used (Gain and François, 2021). SNPs on the assembled chromosomes are shown.

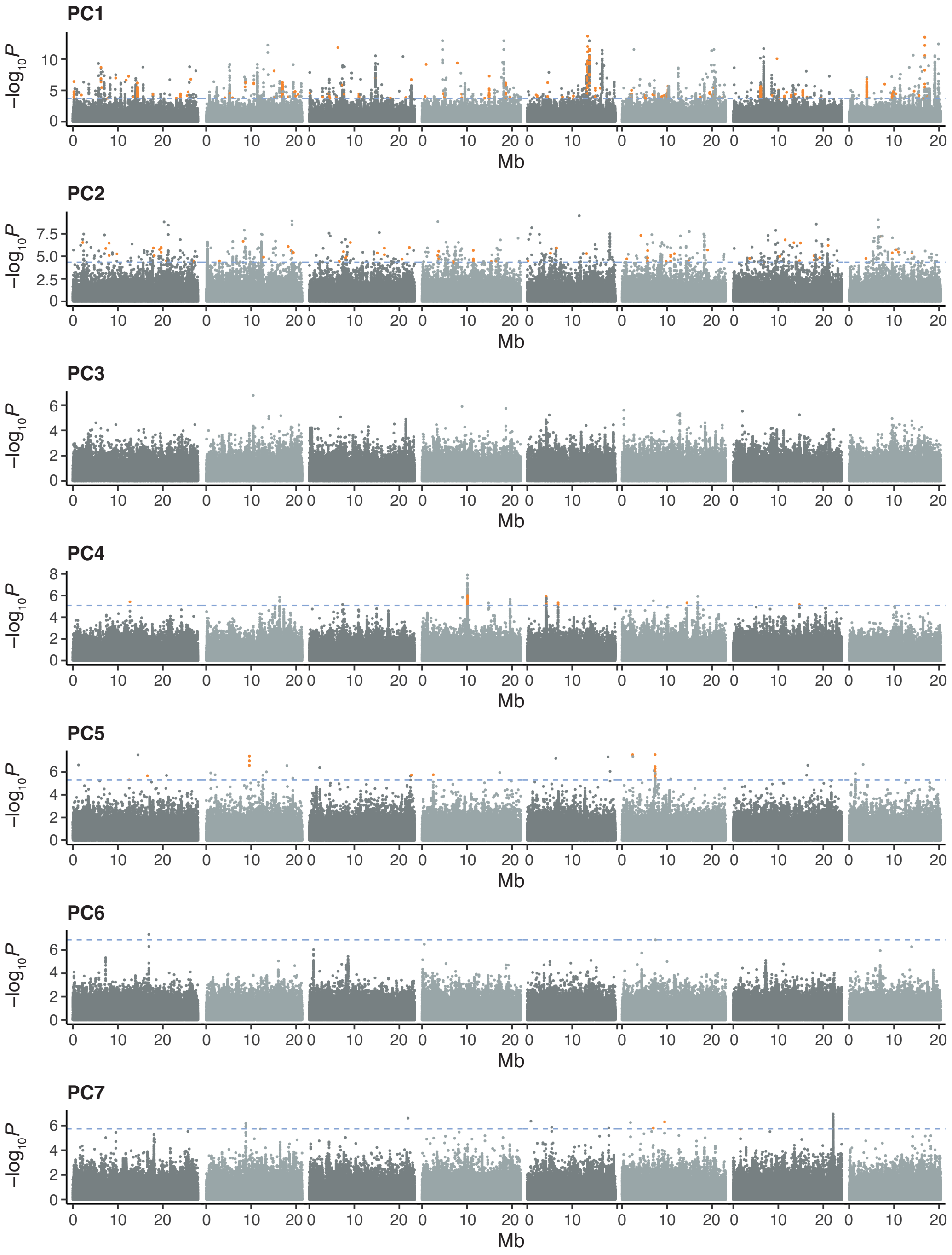


Supplementary Fig. S6. Manhattan plots for principal components of bioclimatic variables. Horizontal dashed lines indicate $P$ = 0.05 after Benjamini–Hochberg false-discovery rate correction. Orange dots indicate SNPs associated significantly with bioclimatic variables and included in the 4-kb windows of selection scans that were significant in more than two combinations of subpopulations. Latent factor mixed models in the R package LEA was used (Gain and François, 2021). SNPs on the assembled chromosomes are shown.

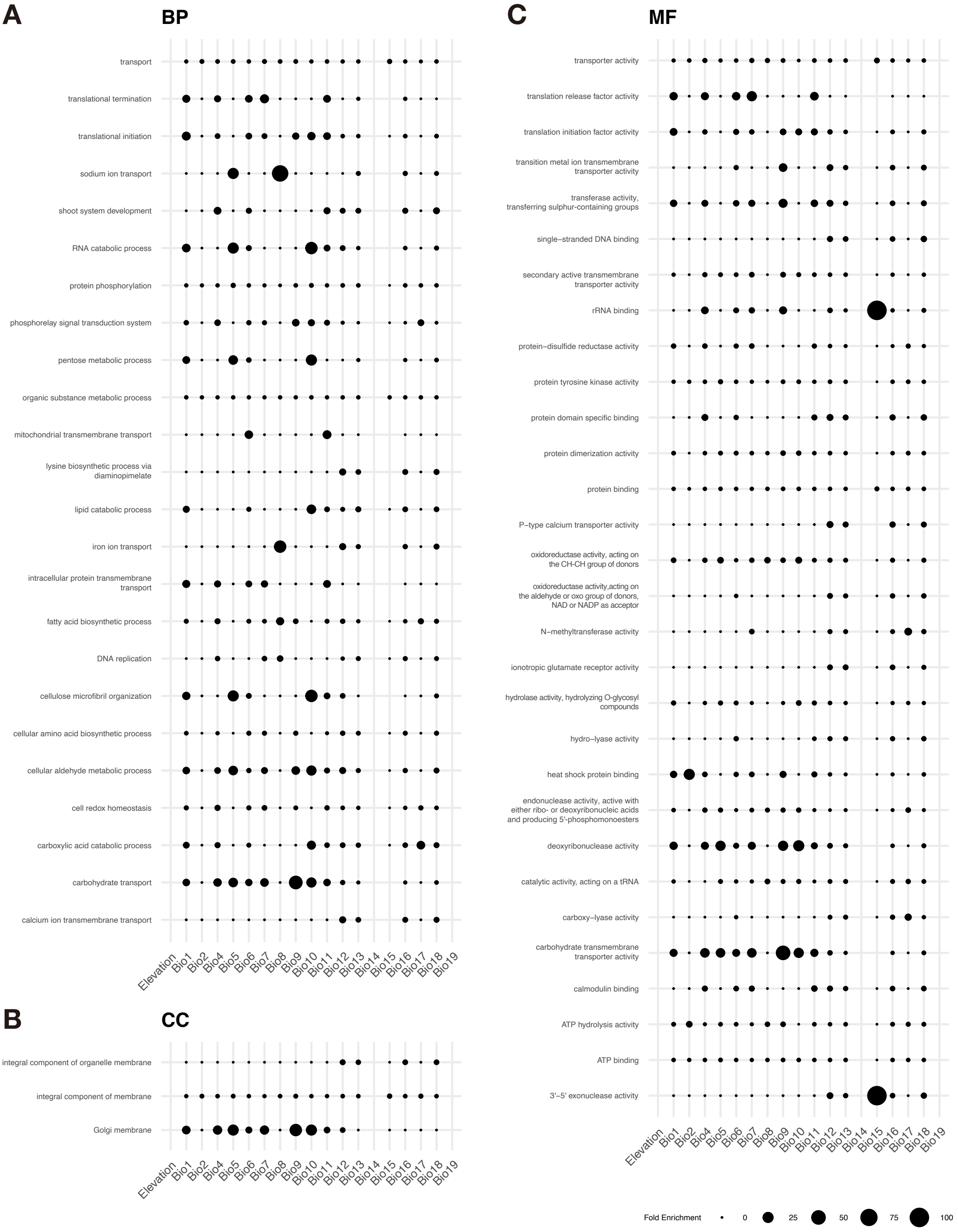


Supplementary Fig. S7. Gene ontology (GO) analysis of climate-associated genes. Enrichments of biological processes (A), cellular components (B), and molecular functions (C) are shown. GO terms with the enrichment of climate-associated genes in at least two bioclimate variables are shown ($P < 0.05$; Fisher's exact test with the “weight01” algorithm in the R package topGO, Alexa et al. 2006). The dot size indicates the fold enrichment.

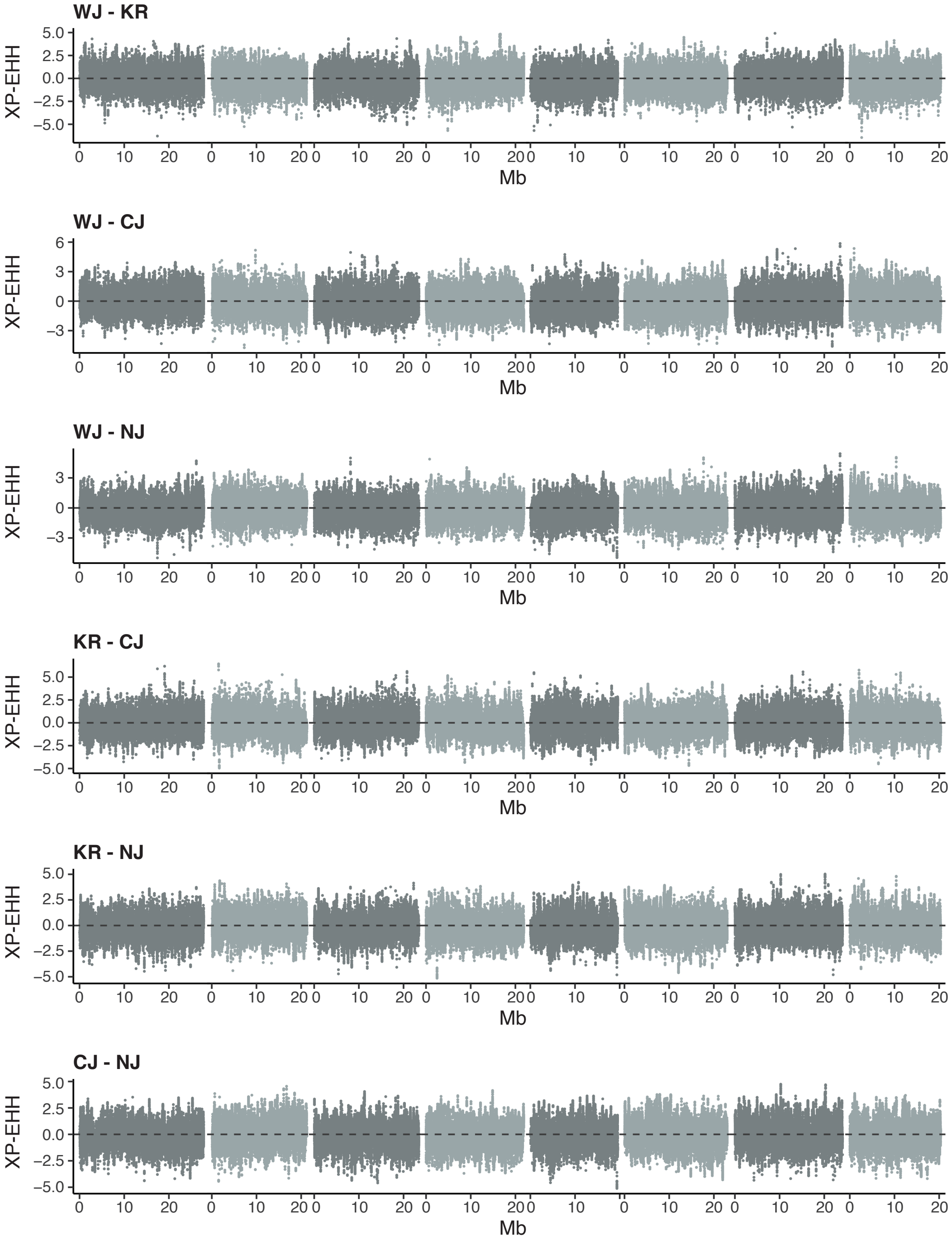


Supplementary Fig. S8. The cross population extended haplotype homozygosity analysis for all the pairwise combinations of subpopulations. The R package rehh v.3.2.2 was used (Gautier et al., 2017). SNPs on the assembled chromosomes were used for analysis.

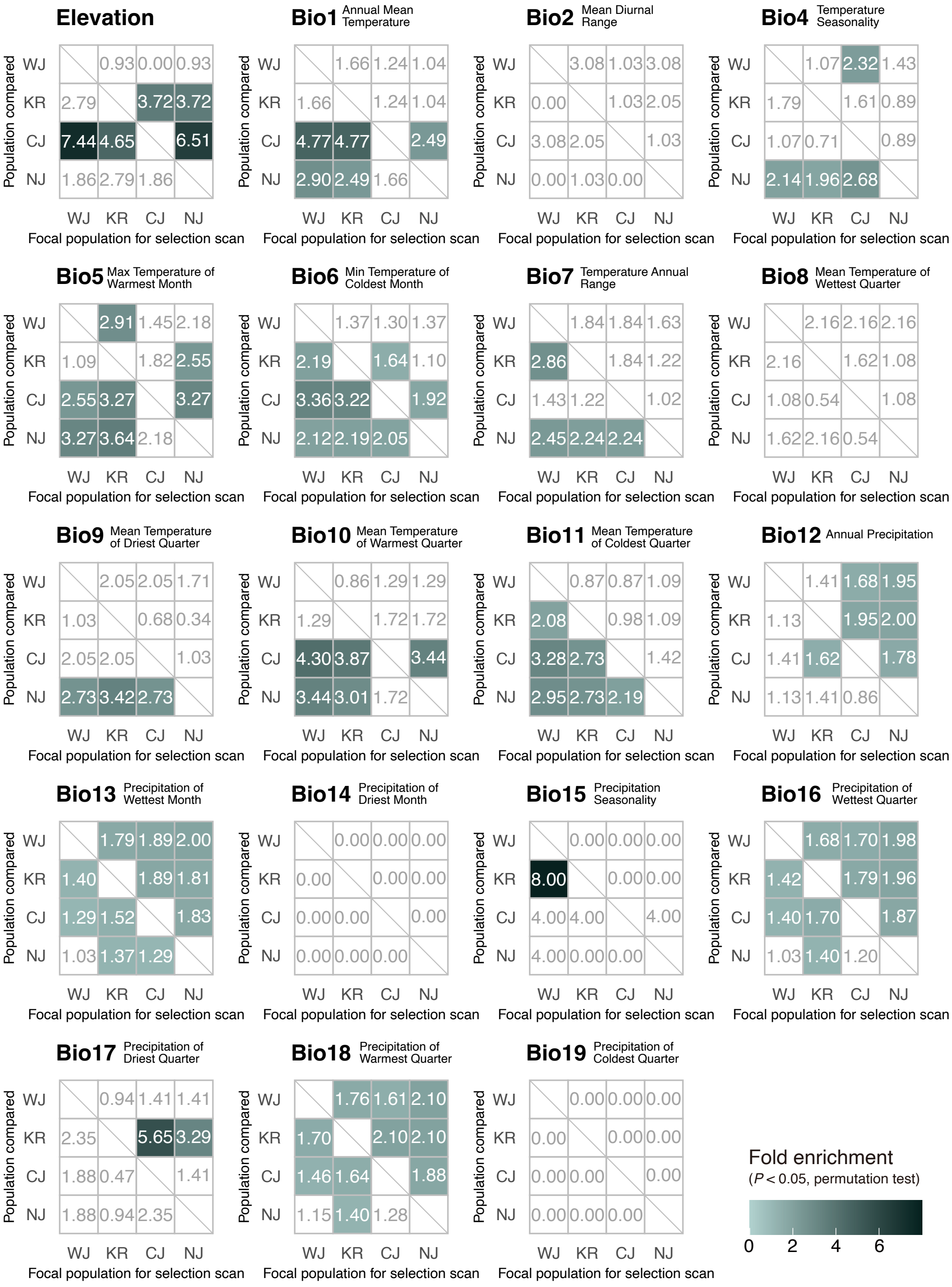


Supplementary Fig. S9. Selection enrichment analysis for all bioclimatic variables. See Fig. 3 for the scheme of the analysis.

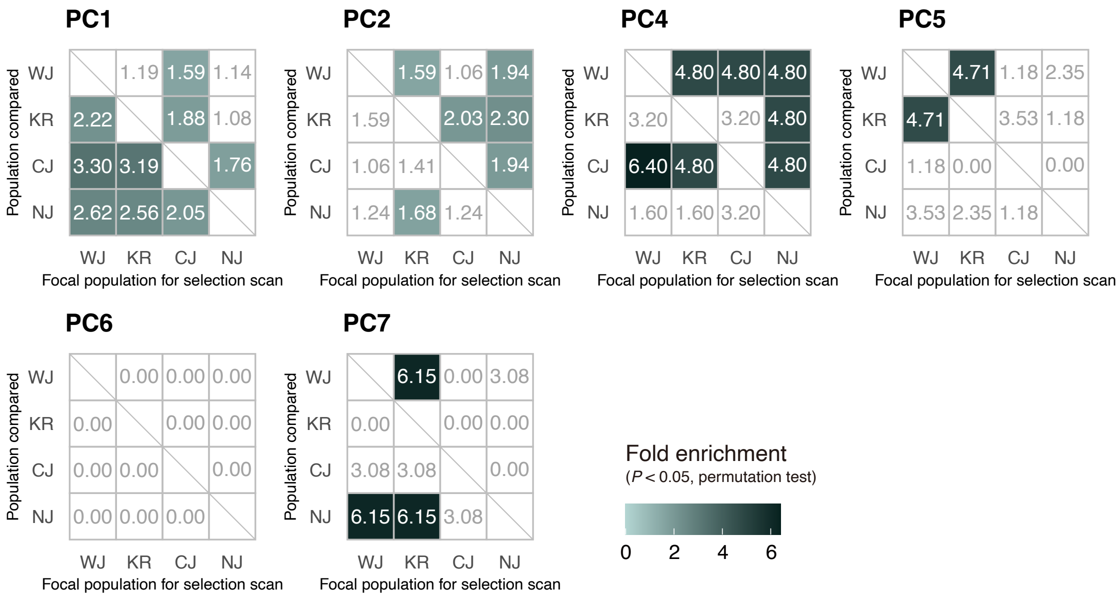


Supplementary Fig. S10. Selection enrichment analysis for principal components of bioclimatic variables. See Fig. 3 for the scheme of the analysis.

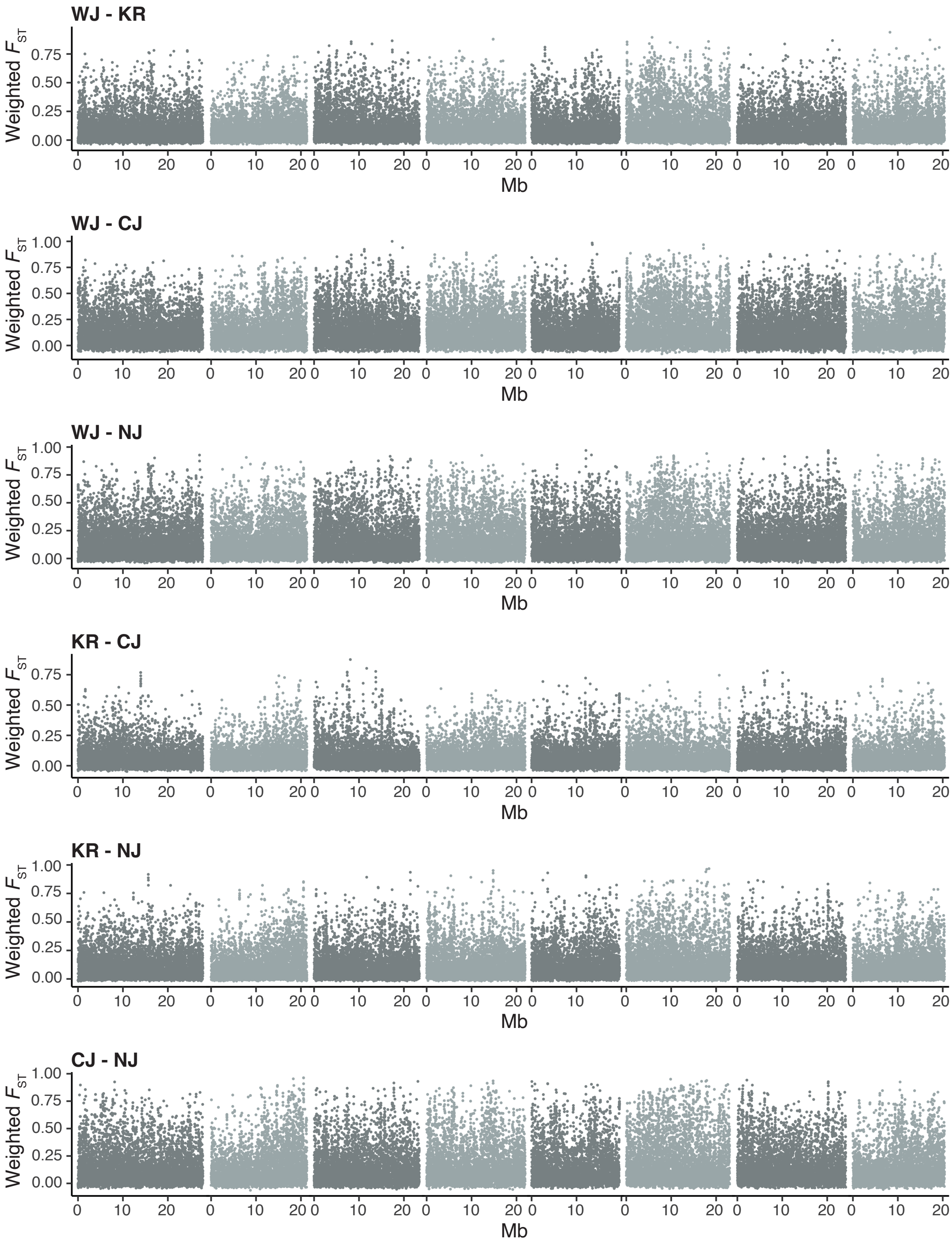


Supplementary Fig. S11. Genome-wide distributions of weighted $F_{ST}$ for pairwise combinations of subpopulations. Weighted $F_{ST}$ in the 4-kb window was calculated using VCFtools v. 0.1.16 (Danecek et al., 2011). SNPs on the assembled chromosomes are shown.

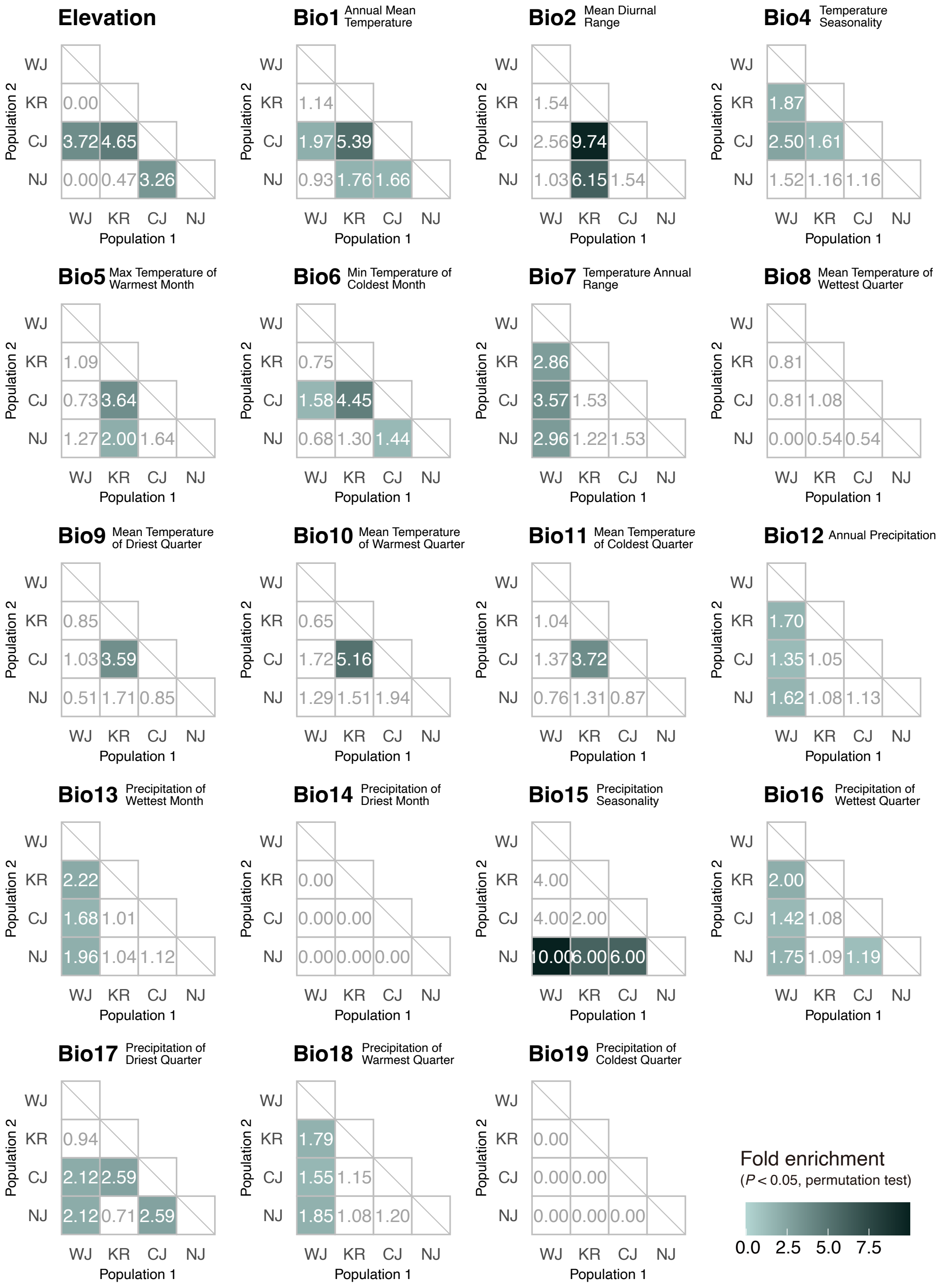


Supplementary Fig. S12. Selection enrichment analysis for all bioclimatic variables, using weighted $F_{ST}$ instead of XP-EHH.

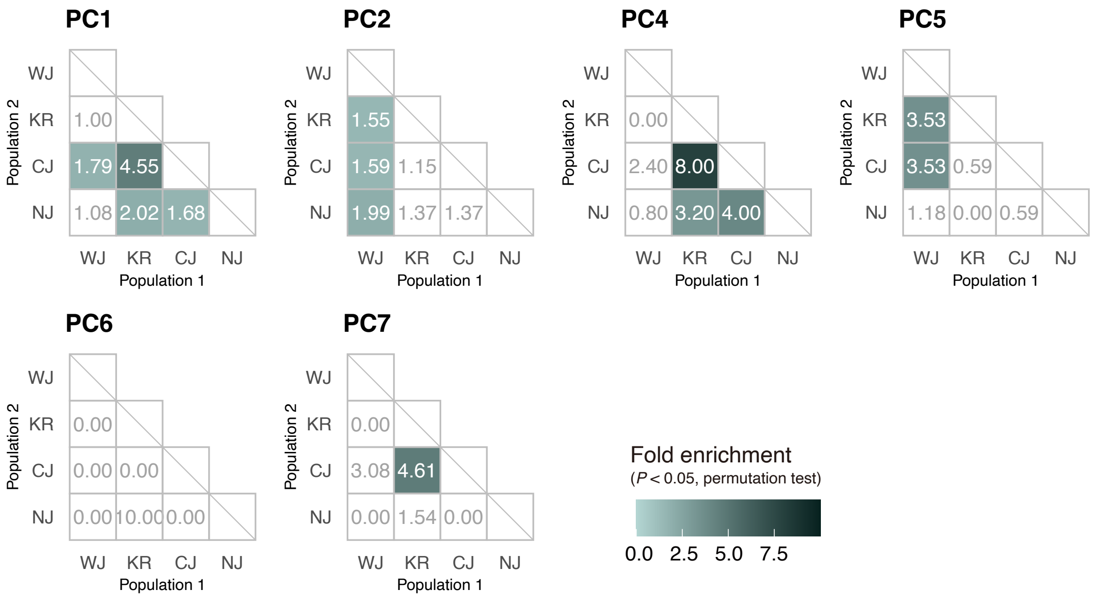


Supplementary Fig. S13. Selection enrichment analysis for principal components of bioclimatic variables, using weighted $F_{ST}$ instead of XP-EHH.

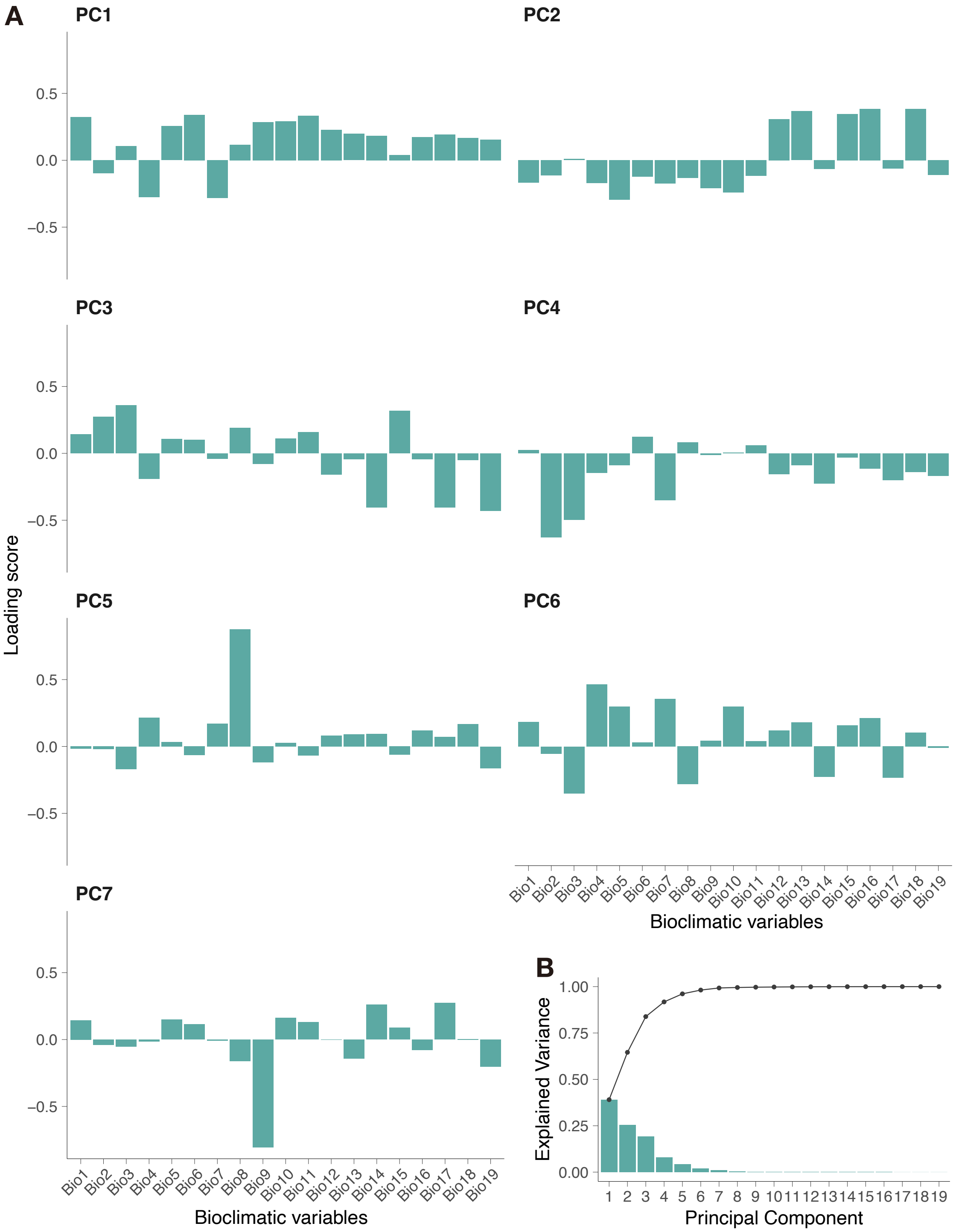


Supplementary Fig. S14. Principal component analysis of bioclimatic variables used in selection scans. (A) Loading scores of each bioclimatic variable for PC1–PC7. (B) Variance explained by each principal component, PC1-PC19 (bar plot). Cumulative explained variance is shown by a line.

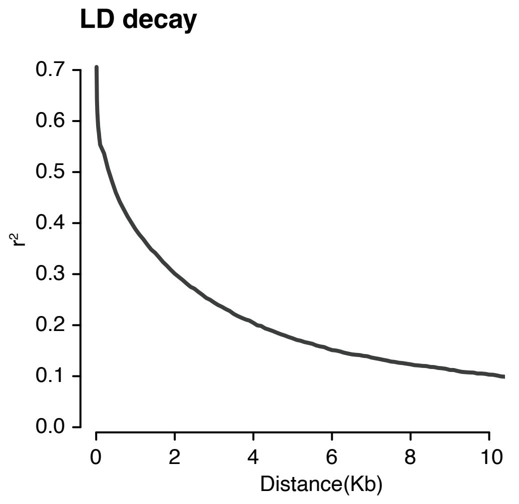


Supplementary Fig. S15. A genome-wide pattern of linkage disequilibrium ($r^2$) decay estimated by PopLDdecay (Zhang et al., 2019).

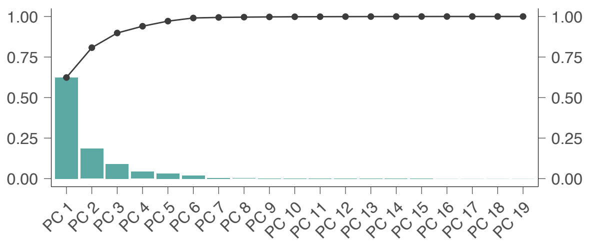


Supplementary Fig. S16. Principal component analysis of bioclimatic variables used in ecological niche modeling. Bar plot indicates variance explained by each principal component. Cumulative explained variance is shown by a line.
